# Polo-like kinase 1 maintains transcription and chromosomal accessibility during mitosis

**DOI:** 10.1101/2025.02.12.637959

**Authors:** Zhouyuan Shen, Kristin Adams, Ryan Moreno, Robert Lera, Emily Kaufman, Jessica D. Lang, Mark Burkard

## Abstract

Transcription persists at low levels in mitotic cells and plays essential roles in mitotic fidelity and chromosomal dynamics. However, the detailed regulatory network of mitotic transcription remains largely unresolved. Here, we report the novel role of Polo-like kinase 1 (Plk1) in maintaining mitotic transcription. Using 5-ethynyl uridine (5-EU) labeling of nascent RNAs, we found that Plk1 inhibition leads to significant downregulation of nascent transcription in prometaphase cells. Chromatin-localized Plk1 activity is required for transcription regulation and mitotic fidelity. Plk1 sustains global chromosomal accessibility in mitosis, especially at promoter and transcription start site (promoter-TSS) regions, facilitating transcription factor binding and ensuring proper transcriptional activity. We identified SMC4, a common subunit of condensin I and II, as a potential Plk1 substrate. Plk1 activity is fundamental to these processes across non-transformed and transformed cell lines, underscoring its critical role in cell cycle regulation. This study elucidates a novel regulatory mechanism of global mitotic transcription, advancing our understanding of cell cycle control.

**Significance Statement:**

- Cells retain a low level of transcription during mitosis, while the regulatory network and specific contributions of mitotic transcription are not well understood.
- We identify Polo-like kinase 1 (Plk1) as a novel regulator of mitotic transcription, crucial for chromosome condensation, genome accessibility, and maintaining mitotic fidelity.
- This study enhances our understanding of Plk1’s multifaceted role in mitotic progression, advancing cell cycle regulation knowledge, and informing new cancer therapies’ development.

## Introduction

Eukaryotic transcription oscillates significantly in a cell cycle-dependent manner, meticulously orchestrated by DNA accessibility, RNA polymerases (RNAPs), and transcription regulators. In mitosis, transcription markedly decreases relative to interphase cells, as first reported in 1962 using nascent RNA labeling in HeLa cells (Prescott & Bender, 1962). This reduced transcription can be explained by key cellular events that render mitotic chromosomes “transcriptionally silent” (Gebara et al., 1997; Gibcus et al., 2018; Gottesfeld & Forbes, 1997; John & Workman, 1998; Martínez-Balbás et al., 1995; Sarge & Park-Sarge, 2009; Segil et al., 1996; Smoyer & Jaspersen, 2014): in early stages of eukaryotic mitosis, the nuclear envelope is disassembled, DNA is condensed into distinct chromosomes, RNAPs (Guo et al., 2014; Liang et al., 2015) and numerous transcription factors (Martínez-Balbás et al., 1995) are phosphorylated, limiting access to the DNA. Recent studies, however, have challenged the view of complete transcriptional silence in mitosis.

For instance, immunofluorescence imaging revealed transcription elongating RNAPII signals at mitotic centromeres, suggesting active transcription of alpha-satellite DNAs (Chan et al., 2012). Pulse-chase labeling with 5-ethynyl uridine (5-EU) (Breinbauer & Köhn, 2003; Jao & Salic, 2008; Kolb et al., 2001; Rostovtsev et al., 2002; Q. Wang et al., 2003) of nascent transcripts in mitotic cells revealed that approximately 25% of expressed genes retain mitotic transcription, albeit at a five-fold reduced level on average compared to interphase (Palozola et al., 2017). Notably, specific genes, accounting for approximately 5% of mitotic transcriptome, exhibited increased expression. This preservation of low-level mitotic transcription implies regulatory mechanisms and functional roles distinct from those well understood during interphase.

Mitotic transcription is not merely a passive residual activity; it may serve distinct regulatory and functional roles. For instance, persistent transcriptional activity at mitotic centromeres maintains centromere integrity and function during cell division (Perea-Resa & Blower, 2018). Indeed, centromeric transcription contributes to the localization of CENP-C (Lyn Chan & Wong, 2012; Rošić et al., 2014), Shugoshin 1 (Liu et al., 2015), Aurora kinase B (Blower, 2016), and ATR (Kabeche et al., 2018), which play key roles in centromere cohesion, kinetochore assembly, and chromosome segregation. Additionally, the upregulation of certain genes during mitosis may be crucial for ensuring the rapid reactivation of transcription programs and timely expression of critical genes upon mitotic exit (Palozola et al., 2017). Mitotically enriched genes are involved in extracellular structure and transcription, with some showing prioritized transcription at mitotic exit (Palozola et al., 2017), suggesting a pivotal role in maintaining cell differentiation and identity. The differential expression patterns observed during mitosis and mitotic exit indicate a tightly regulated process that ensures cellular homeostasis and functional continuity through the cell cycle. Despite these significant advances, questions remain regarding detailed function of mitotic enriched transcripts and potential roles of the observed basal transcriptional activity on chromosomes arms.

The persistent mitotic transcription at different mitotic stages requires dynamic arrangement of transcription machineries. Consistent with transcription retention, mitotic cells maintain moderate genome-wide accessibility, as measured by assay for transposase-accessible chromatin sequencing (ATAC-seq) (Hsiung et al., 2014; Teves et al., 2016; Yu et al., 2023). This accessibility allows transcriptional machinery to access necessary DNA regions, even during the highly condensed state of mitosis. Mitotic transcription at centromeres and chromosomal arms is RNAP II-dependent, as selective inhibition of RNAP II abolishes active transcription signals (Chan et al., 2012; Palozola et al., 2017). Aberrant retention of RNAP II on mitotic chromosomes causes delays in cell cycle progression and improper transcription reactivation post-mitosis (Perea-Resa et al., 2020; Wiegard et al., 2021), underscoring the delicate balance needed for proper mitotic progression. These findings highlight the potential consequences of transcriptional dysregulation during cell division and also the necessity of understanding the mechanisms regulating mitotic transcription.

Polo-like kinase 1 (Plk1) is a serine/threonine kinase that plays crucial roles in multiple stages of the cell cycle. The expression levels of PLK1 increase during the G2 phase and peak during the M phase. Plk1 exhibits dynamic subcellular localization throughout mitosis, correlating with its functions at different mitotic stages. During interphase, Plk1 localizes to centrosomes, where it promotes centrosome separation and maturation (Casenghi et al., 2005; Conduit et al., 2014; Haren et al., 2009; Lee & Rhee, 2011; Oshimori et al., 2006; Tsou et al., 2009). In prophase, Plk1 regulates nuclear envelope breakdown (Tsou et al., 2009) and chromosome arm resolution (Hauf et al., 2005); its localization and activity are required at chromosome arms. During prometaphase, centrosome-localized Plk1 controls bipolar spindle assembly (Oshimori et al., 2006), while a portion of the kinase is recruited by Bub1 and CENPU to kinetochores (Q. Chen et al., 2021; Nguyen et al., 2021; Singh et al., 2021), where it facilitates proper kinetochore–microtubule (KT–MT) attachments. As cells progress to anaphase, Plk1 dissipates from kinetochores and relocates to the central spindle and spindle midbody, where it coordinates cytokinesis and cell abscission (Beck et al., 2013; Burkard et al., 2007, 2009; Golsteyn et al., 1995). Inhibition of Plk1 activity generates a spectrum of mitotic phenotypes, depending on the timing and level of inhibition. Aberrant Plk1 activity can result in cell cycle progression delays and improper mitotic exit, underscoring the delicate balance required for proper cell division. These observations highlight the importance of Plk1 in mitotic regulation and the need for precise control of Plk1 activity. Continued study of Plk1 and its interactions is essential for a comprehensive understanding of cell cycle regulation and genomic stability maintenance.

Polo-like kinase 1 recognizes its target proteins through the interaction of its Polo-box domain (PBD) with phosphopeptides (Jang et al., 2002; Seki et al., 2008). Although Plk1 does not directly interact with DNA, evidence suggests that it plays a critical role in transcription regulation by modulating the chromatin landscape, transcription factors, and transcriptional machinery. For instance, Plk1 directly interacts and phosphorylates histone demethylase LSD1 at Ser-126, promoting its release from chromatin during mitosis (Peng et al., 2017), which affects histone methylation levels and likely chromatin accessibility. Additionally, Plk1 phosphorylates and activates transcription factors that are crucial for the transcription of genes involved in cell cycle progression and mitotic spindle assembly. During the G2/M transition, Plk1 binds to and phosphorylates FoxM1, enhancing its activity and increasing the expression of a cluster of G2/M phase target genes, including PLK1, thus establishing a positive feedback loop (Fu et al., 2008; Laoukili et al., 2005; Thiru et al., 2014; I.-C. Wang et al., 2005; Wonsey & Follettie, 2005). However, current studies have not differentiated the activity and function of Plk1 across different cell cycle stages, leaving unanswered questions regarding the precise timing and subcellular localization of Plk1’s functions.

Plk1 licenses new CENP-A deposition at centromeres during the early G1 phase of the cell cycle, which is essential for maintaining centromere identity (Conti et al., 2024). Centromeric transcription plays a significant role in CENP-A loading, with transcriptional inhibition resulting in decreased CENP-A at centromeres (Bergmann et al., 2012; Cardinale et al., 2009; Catania et al., 2015; Choi et al., 2011; Nakano et al., 2008; Zhu et al., 2018). Transcription by RNAP II and concomitant chromatin remodeling are required to mediate the transition of CENP-A from a chromatin-associated state to a stably incorporated form (Bobkov et al., 2018). During mitosis, partial inhibition of Plk1 leads to a higher rate of chromosome missegregation during anaphase (Addis Jones et al., 2019; Lera et al., 2016, 2019), whereas transcription inhibition is also associated with increased mitotic errors (Liang et al., 2015; Liu et al., 2015; Lyn Chan & Wong, 2012). These findings suggest a potential link between Plk1 and transcription at centromeres and other chromosomal regions during mitosis; however, direct studies examining this relationship are lacking.

Here we report the discovery that Polo-like kinase 1 (Plk1) maintains cellular transcription during mitosis. We find that inhibiting Plk1 activity markedly decreases mitotic transcription. Furthermore, Plk1 limits chromosome condensation, maintaining chromosomal accessibility and thereby allows transcription machinery to access mitotic DNA. Using chemical genetic tools, we find that Plk1 acts directly on chromatin to sustain chromatin accessibility and transcription, and this activity is required for fidelity of chromosome segregation. This effect is partially mediated through the phosphorylation of Structural maintenance of chromosomes protein 4 (SMC4), a key condensin component. These data demonstrate a molecular mechanism by which Plk1 activity regulates the chromosomal landscape and transcriptional activities during mitosis.

## Results

### Plk1 Activity Maintains Mitotic Transcription

To investigate mitotic transcription during normal cell mitosis, we use hTERT-immortalized human retinal pigment epithelial-1 (RPE1) cells. After treating with Eg-5 inhibitor S-trityl-l-cysteine (STLC) (DeBonis et al., 2004; Wu et al., 2018) for 2 hours, the cells were arrested at prometaphase due to the failure of spindle bipolarization and subsequent collapse into monopolar structures. 5-ethynyl uridine (5-EU) was added during the 2-hour drug treatment, labeling all nascent transcripts synthesized within this period. Click reaction visualization of 5-EU-labeled nascent transcripts in prometaphase cells revealed a robust fluorescent staining pattern permeating the entire cell, while the no 5-EU addition group showed a complete loss of fluorescent signal (Fig. S1A). The signal overlapped with DNA and displayed a distinct enrichment outlining the chromosomes, indicating that nascent transcription persists on mitotic chromosomes. Thus, we successfully visualized transcription in mitotic cells.

We next examined the effect of Plk1 activity on mitotic transcription. Compared to the 5-EU incorporation pattern in the STLC-only group, the addition of the Plk1 inhibitor BI-2536 (Lénárt et al., 2007) alongside STLC induced a similar chromosome arrangement pattern but significantly reduced nascent transcription signals in prometaphase cells (Fig. 1A-B). However, it remained unclear whether these visualized nascent transcripts were synthesized before or after the cells entered mitosis during synchronization and labeling using this method. Moreover, given that Plk1 is activated in late G2 phase and peaks in activity during mitosis (Archambault & Glover, 2009), we sought to distinguish the effect of Plk1 activity on G2 or M phase by further segregating cells at different cell cycle stages. Cells were synchronized through thymidine block for S phase enrichment, then released into fresh media to proceed into G2 and mitosis. Mitotic cells were collected by shake-off and incubated in 5-EU containing media simultaneously with their corresponding interphase cells. Chromosome spread staining enabled visualization of chromosome-bound nascent RNAs (Fig. 1C). Interphase and mitotic spreads were individually quantified and compared with or without Plk1 inhibition, ensuring that the nascent transcripts were labeled and were directly influenced by Plk1 activity during the respective cell cycle stage. While a five-fold reduction in transcription was observed between mitotic and interphase cells, consistent with previous studies (Palozola et al., 2017), Plk1 inhibition significantly decreased 5-EU staining in mitotic cells but not in interphase cells. (Fig. 1D-E). This finding indicates a mitotic-specific effect of Plk1 activity on transcription regulation. The reliance on Plk1 activity for mitotic transcription was also observed in transformed cell lines including HeLa and Cal51 (Fig. 1F-G, Fig. S1B-D), suggesting a conserved basic cellular regulatory mechanism.

**Figure 1.**
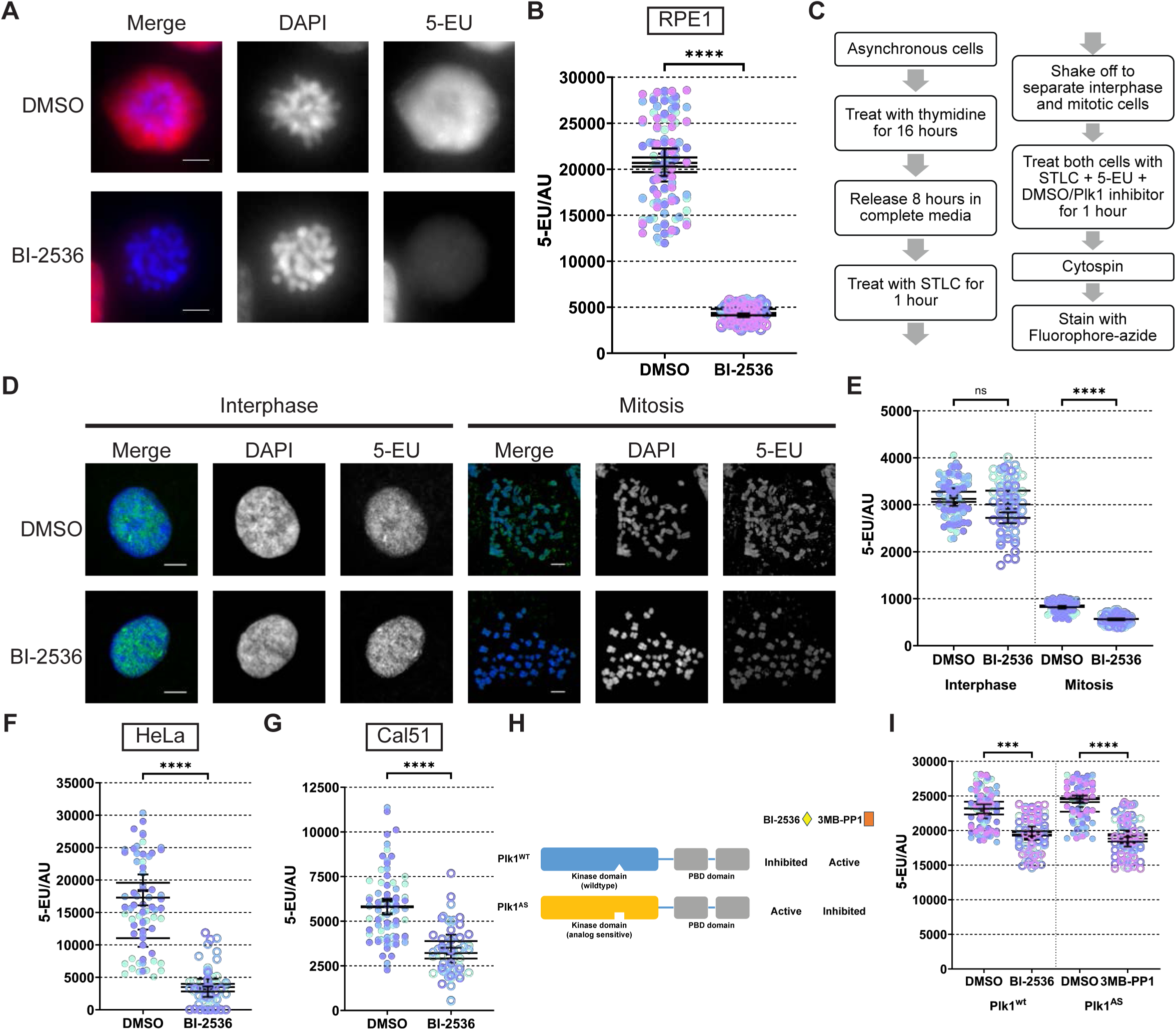
Plk1 activity maintains transcription during mitosis. (A) Representative images of 5-EU staining corresponding to nascent transcription in prometaphase RPE1 cells. DMSO or final concentration of 200 nM Plk1 inhibitor BI-2536 were added in 1 mM 5-EU and 5 µM STLC containing media for 2 hours. (B) Quantification of chromosomal 5-EU fluorescent signal in prometaphase RPE1 cells showed in (A). (C) Schematic of separating mitotic and interphase cells for 5-EU incorporation and nascent transcript staining. (D) Representative images of RPE1 interphase and mitotic cell spreads after treatment in (C). (E) Quantification of chromosomal 5-EU fluorescent signal in spreads of RPE1 interphase and mitotic cells showed in (C). (F) Quantification of chromosomal 5-EU fluorescent signal in prometaphase HeLa cells treated with DMSO or 200 nM BI-2536 for 2 hours. (G) Quantification of chromosomal 5-EU fluorescent signal in prometaphase Cal-51 cells treated with DMSO or 200 nM BI-2536 for 2 hours. (H) Design of analog sensitive (AS) Plk1 system and the inhibition specificity in RPE1 Plk1^-/-^ EGFP-Plk1^WT^ (Plk1^WT^) and RPE1 Plk1^-/-^ EGFP-Plk1^AS^ (Plk1^AS^) cells. (I) Quantification of chromosomal 5-EU fluorescent signal in prometaphase Plk1^WT^ and Plk1^AS^ cells treated with DMSO or corresponding Plk1 inhibitors for 2 hours. For microscopic images, scale bar: 5 µm. For 5-EU quantification, each dot represents a cell, and different colors represent different biological replicates. 3 or more biological replicates were examined.

The chemical BI-2536 efficiently inhibits Plk1 activity at nanomolar concentrations and also affects BRD4, a transcription regulator (Ciceri et al., 2014; Ember et al., 2014). To eliminate any off-target effects of BI-2536 on transcription repression, we employed the analog-sensitive Plk1 (Plk1^AS^) system (Fig. 1H). Plk1^AS^ harbors mutations in two gatekeeper residues which prevent inhibition by BI-2536 but sensitize the kinase to the bulky ATP analog 3MB-PP1 (Burkard et al., 2007, 2012; Lera & Burkard, 2012), demonstrating kinase function specificity. In mitotic RPE1 Plk1^-/-^ cells expressing EGFP-tagged wild-type Plk1 (Plk1^WT^) or Plk1^AS^, specific inhibition of Plk1 activity resulted in similar transcription repression (Fig. 1I), confirming that Plk1 enzymatic activity is required to maintain mitotic transcription. Quantification of nascent transcripts retained on mitotic chromosomes in Plk1^AS^ cells using chromosome spreads revealed a downregulation following Plk1 inhibition (Fig. S1E-F). Finally, transcription downregulation was observed in RPE1 cells using a structurally distinct Plk1 inhibitor, GSK461364 (Gilmartin et al., 2009) (Fig. S1G). Therefore, we concluded that Plk1 activity is essential for maintaining transcription levels in the later stages of mitosis.

### Chromosomal Plk1 Activity is Required for Mitotic Transcription and Chromosomal Segregation Fidelity

In light of the discovery of Plk1’s regulatory role in mitotic transcription, we investigated the effects on mitotic transcription, alongside the detailed mechanisms and roles Plk1 plays during this process. Previous studies suggested that mitotic transcription is essential for maintaining mitotic fidelity. Indeed, in our experiments, treating asynchronous RPE1 cells with the RNAP II inhibitor α-amanitin for 5 hours (Kume et al., 2016; Zhao et al., 2006), resulted in a significant increase in missegregation rates during anaphase compared to the control group (Fig. 2A-B), consistent with earlier reports (Chan et al., 2012; Liang et al., 2015; Liu et al., 2015). Plk1^AS^ cells exhibited a similar effect following α-amanitin treatment (Fig. 2C). To validate the results of pharmacological inhibition, we acutely depleted the endogenous RNAP II subunit POLR2A in an Auxin-inducible degron cell line (DLD1 tet-OsTIR1 mClover-mAID-POLR2A, Nagashima et al., 2019, short for POLR2A-AID). Addition of IAA diminished mClover signal corresponding to POLR2A protein in live cell imaging (Fig. S2A). Reduced RNAP II levels led to a significant increase in mitotic errors, including lagging chromosomes and chromosome bridging (Fig. 2D-E). Notably, partial Plk1 inhibition, as reported by our group and others (Addis Jones et al., 2019; Lera et al., 2016, 2019), also resulted in higher missegregation rates during anaphase (Fig. 2B, 2C, and 2E). Thus, it is possible that Plk1 activity maintains mitotic chromosomal segregation fidelity by regulating mitotic transcription, although it remains formally possible that some suppressed transcripts in these experiments are not regulated by Plk1.

**Figure 2.**
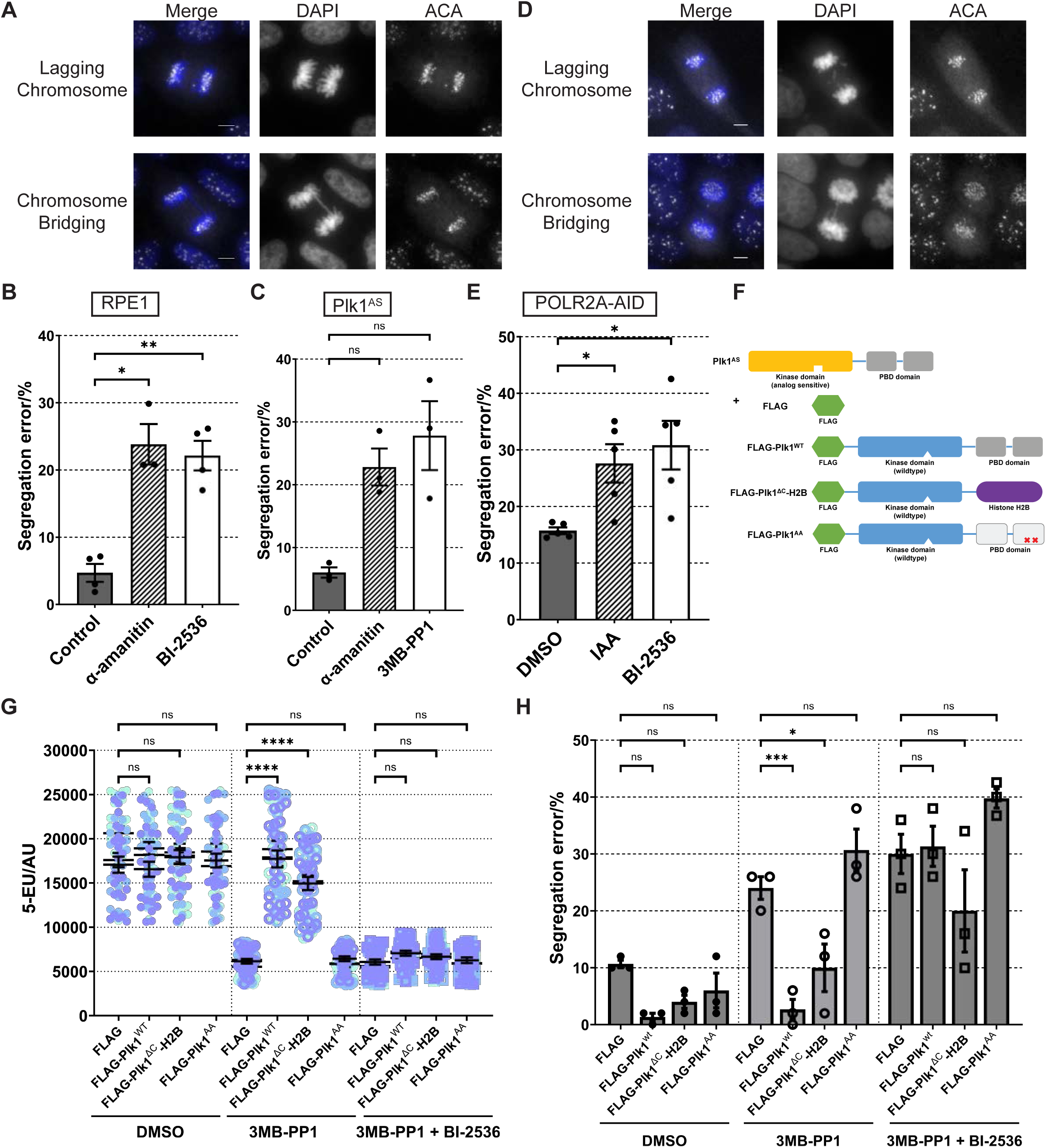
Chromosomal Plk1 Activity is Required for Mitotic Transcription and Chromosomal Segregation Fidelity. (A) Representative images of anaphase RPE1 cells with erroneous chromosome segregation after 5-hour treatment of final concentration of 50 µg/mL α-amanitin. (B) Quantification of erroneous chromosome segregation in anaphase RPE1 cells treated with DMSO, 50 µg/mL α-amanitin, or 10 nM BI-2536 for 5 hours. (C) Quantification of erroneous chromosome segregation in anaphase Plk1^AS^ cells treated with DMSO, 50 µg/mL α-amanitin, or 200 nM 3MB-PP1 for 5 hours. (D) Representative images of anaphase DLD1 tet-OsTIR1 mClover-mAID-POLR2A (POLR2A-AID) cells with erroneous chromosome segregation after 4-hour 0.5 mM IAA treatment. (E) Quantification of erroneous chromosome segregation in anaphase POLR2A-AID cells pretreated for 23-hour 1 μg/mL doxycycline, and incubate in DMSO, 0.5 mM IAA or 10 nM BI-2536 containing media for 4 hours. (F) Illustration of the design of subcellular localization-specific Plk1 activity rescue system in Plk1^AS^ lines. (G) Quantification of prometaphase chromosomal 5-EU fluorescent signal in indicated cell lines treated with DMSO, 10 µM 3MB-PP1, or 10 µM 3MB-PP1+200 nM BI-2536 for 2 hours with the presence of STLC and 5-EU. (H) Quantification of erroneous anaphase chromosome segregation in indicated cell lines treated with DMSO, 200 nM 3MB-PP1, or 200 nM 3MB-PP1+200 nM BI-2536 for 4 hours. For microscopic images, scale bar: 5 µm. For 5-EU quantification, each dot represents a cell, and different colors represent different biological replicates. For mitotic error quantification, each dot represents a biological replicate (>50 anaphase cells were analyzed). 3 or more biological replicates were examined.

Active Plk1 kinase has multiple subcellular locations throughout mitosis. To ascertain whether chromatin-based Plk1 activity regulates mitotic transcription, we utilized a Plk1 activity rescue system developed in our lab (Lera et al., 2016, 2019) (Fig. 2F). Briefly, we introduced FLAG-tagged Plk1 variants in Plk1^AS^ cells. In addition to Plk1^WT^ and Plk1^AA^ (mutating H538A, K540A within the PBD, blocking Plk1 binding specificity (Burkard et al., 2009; Elia et al., 2003)), we replaced the C-terminal Polo-box domain (PBD) of wild-type Plk1 protein (C-terminal truncated, Plk1^ΔC^) with histone H2B (FLAG-H2B-Plk1^ΔC^). In FLAG-H2B-Plk1^ΔC^ cells, Plk1 kinase activity is expressed and is restricted to chromosomes, as demonstrated by FLAG-tag chromosomal localization (Fig. S2B). When global Plk1^AS^ activity was inhibited by 3MB-PP1, this fusion protein provided Plk1 enzymatic activity at specific locations, achieving partial rescue for local Plk1 requirements. In Plk1^AS^ cells, the addition of 3MB-PP1 led to reduction of mitotic transcription at prometaphase (Fig. 1H, Fig. 2G) and increase of chromosome missegregation rates in anaphase cells (Fig. 2C&H). Additional expression of H2B-Plk1^ΔC^ partially rescued prometaphase transcription and anaphase segregation fidelity phenotypes (Fig. 2G-H). Addition of BI-2536 to block the kinase function of Plk1^ΔC^ abolished all the effects, confirming the requirement of chromosomal localized Plk1 activity. As controls, Plk1^WT^ fully rescued all mentioned phenotypes, while Plk1^AA^ showed no difference compared to Plk1^AS^ cells. Therefore, chromosomal localization of Plk1 activity in mitotic cells is critical for transcription and anaphase chromosomal segregation.

In conclusion, Plk1 activity on chromosomes is vital for maintaining mitotic transcription and segregation fidelity.

### Plk1 Regulates Mitotic Transcription Through Chromosome Condensation

Next, we investigated the mechanism by which Plk1 affects mitotic transcription. We demonstrated that its chromosomal localization of kinase activity is necessary (Fig. 2) although Plk1 is not known to directly bind DNA. We raised the possibility that Plk1 regulates mitotic transcription by altering the chromosomal landscape.

Inhibition of Plk1 activity results in centrosome separation failure and bipolar spindle formation, which causes cells to arrest with monopolar spindles in prometaphase (Barr et al., 2004; Kishi et al., 2009; Oshimori et al., 2006). Notably, we observed increased chromosome condensation in these Plk1-inhibited cells (Fig. S3A). To control the degree of chromosomal condensation affected by mitotic timing, we used STLC to arrest cells at a similar prometaphase-like stage (Fig. 3A). The addition of BI-2536 did not alter chromosome arrangement in those cells, but visibly increased chromosome density compared to the DMSO control (Fig. 3A).

**Figure 3.**
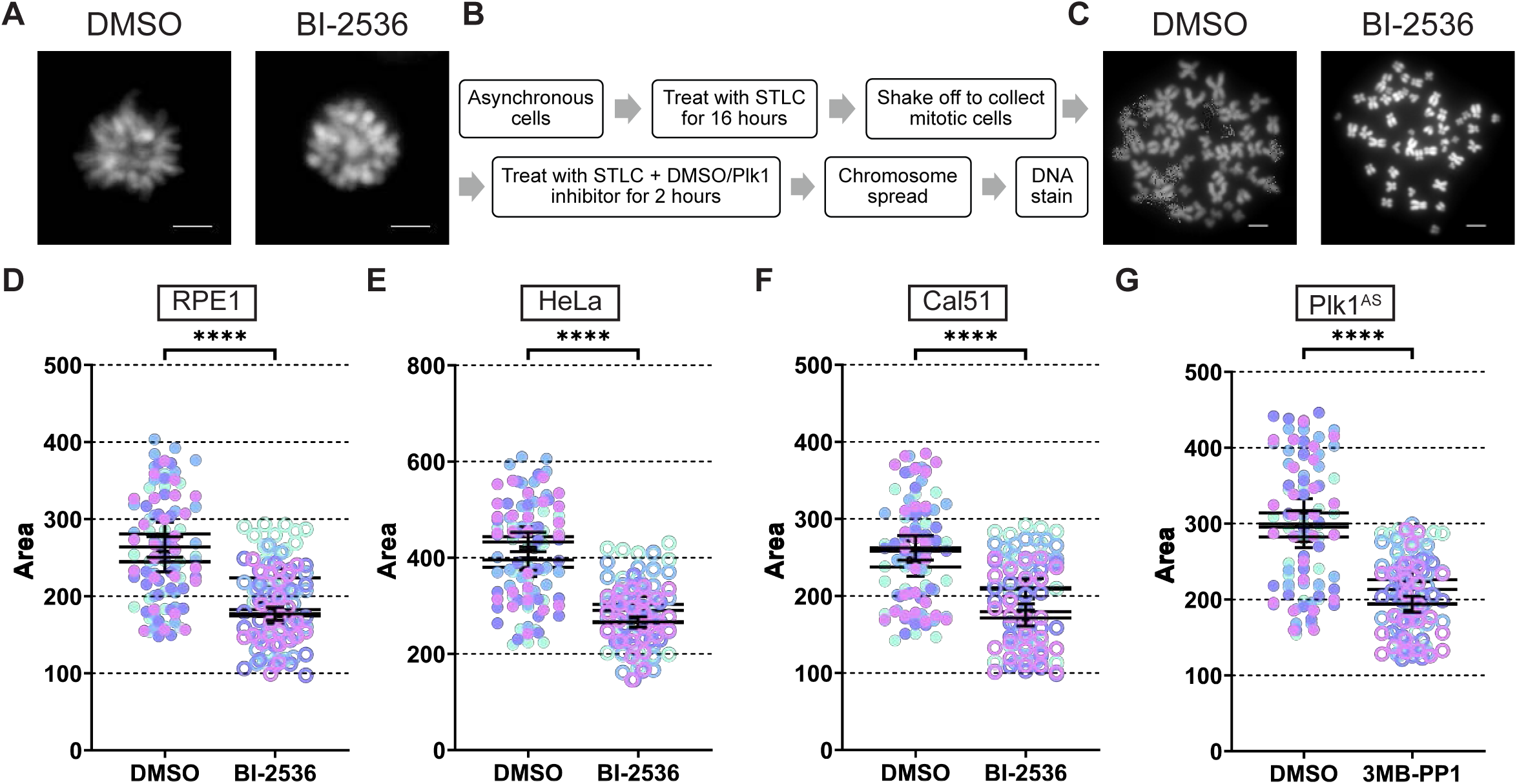
Plk1 activity restricts chromosome condensation levels in prometaphase. (A) Representative images of chromosomal arrangement in STLC synchronized RPE1 prometaphase cells treated with DMSO or 200 nM BI-2536 for 2 hours. (B) Schematic of collecting, treating and processing prometaphase chromosome spreads. (C) Representative images of RPE1 prometaphase chromosomal spreads treated with DMSO or 200 nM BI-2536 for 2 hours. (D) Quantification of chromosomal area in chromosomal spreads in RPE1 prometaphase cells treated with DMSO or 200 nM BI-2536 for 2 hours. (E) Quantification of chromosomal area in chromosomal spreads in HeLa prometaphase cells treated with DMSO or 200 nM BI-2536 for 2 hours. (F) Quantification of chromosomal area in chromosomal spreads in Cal-51 prometaphase cells treated with DMSO or 200 nM BI-2536 for 2 hours. (G) Quantification of chromosomal area in chromosomal spreads in Plk1^WT^ and Plk1^AS^ cells treated with DMSO or corresponding Plk1 inhibitors for 2 hours. For microscopic images, scale bar: 5 µm. For chromosomal condensation quantification, each dot represents a cell, and different colors represent different biological replicates. 3 or more biological replicates were examined.

During mitotic prophase, the kinases Plk1, Aurora B, and Cdk1 phosphorylate the cohesin-associated proteins sororin and SA2 to facilitate cohesin removal from chromosome arms via the “prophase pathway” (Haarhuis et al., 2014). Inhibition of Plk1 results in unresolved sister chromatid arms in post-prophase cells (Haarhuis et al., 2014; Hauf et al., 2005; Sumara et al., 2002). However, we don’t think this role of Plk1 fully explains the over-condensation phenotype we observed. When we collected STLC-synchronized prometaphase cells, treated them with Plk1 inhibitors for 2 hours, and performed chromosome spreads (Fig. 3B), we noticed an over-condensation effect compared to control cells (Fig. 3C). These cells were collected after the cohesin removal step, and we observed fully resolved chromosome arms in mitotic spreads with over-condensed chromosomes. It suggested that despite successful cohesin removal, Plk1 activity inhibition caused chromosome over-condensation. In these cells, Plk1 activity is sufficient for cohesin removal during prophase but is insufficient to maintain proper chromosome condensation levels when arrested in prometaphase.

For quantitative analysis, we measured the DNA area in chromosomal spreads. Inhibition of Plk1 activity in STLC-arrested RPE1 cells decreased chromosome area, confirming Plk1’s role in restraining chromosome condensation (Fig. 3D). This observation extends to HeLa and Cal51 cells (Fig. 3E-F), indicating a common mechanism in both non-transformed and transformed cells. Over-condensation was observed in both Plk1^WT^ cells treated with BI-2536 and Plk1^AS^ cells treated with 3MB-PP1 (Fig. 3G), confirming the specificity of Plk1-mediated condensation regulation. Treatment with another structurally distinct Plk1 inhibitor, GSK461364, yielded similar results in RPE1 cells (Fig. S3B).

These findings suggest that Plk1 activity is essential for regulating chromosome condensation, preventing over-condensation in later mitosis. They also provide a mechanistic explanation for how Plk1 regulates mitotic transcription.

### Plk1 Activity Regulates Chromosomal Accessibility in Mitosis

Chromosomal condensation abolishes most long-range interactions between distant chromosome regions as cells enter mitosis. However, recent studies have found that the local chromosomal landscape is largely preserved in mitotic chromosomes (D. Chen et al., 2005; Hsiung et al., 2015; Ou et al., 2017; Teves et al., 2016). Previous ATACseq results in mouse embryonic stem cells showed that Tn5 transposase accesses mitotic chromosomes as efficiently as interphase chromatin, generating nearly identical integration patterns and sequencing read counts in ATAC-seq profiles (Teves et al., 2016). Therefore, the genome retains global chromosome accessibility and open chromatin status during mitosis. However, the detailed regulatory mechanisms of mitotic accessibility are yet to be uncovered, and the modulation of local accessibility in mitotic chromosomes remains understudied. Considering the effect of Plk1 activity on mitotic chromosome condensation, we evaluate changes in chromosomal accessibility in mitotic cells influenced by Plk1 activity. We hypothesized that Plk1 inhibition, which leads to chromosome over-condensation, would reduce open chromatin levels in mitotic cells.

We performed bulk ATAC-seq in mitotic cells to test this hypothesis. Prometaphase RPE1 Plk1^AS^ cells were synchronized and collected by mitotic shake-off. Cells were then treated with DMSO or the Plk1 inhibitor 3MB-PP1 for 2 hours before ATAC library preparation (Fig. 4A). We observed a similar pattern of fragment length distribution of ATAC-seq reads in both groups (Fig. S4A), consistent with the mitotic cell population from previous study (Teves et al., 2016). Among four biological replicates, consistent enrichment of peaks was observed in both groups (Fig. S4B). Consensus peaks were mapped onto chromosomes in both mitotic populations, suggesting the presence of open chromatin regions in prometaphase cells (Fig. 4B). Peaks were found on every chromosome arm (Fig. S4C).

**Figure 4.**
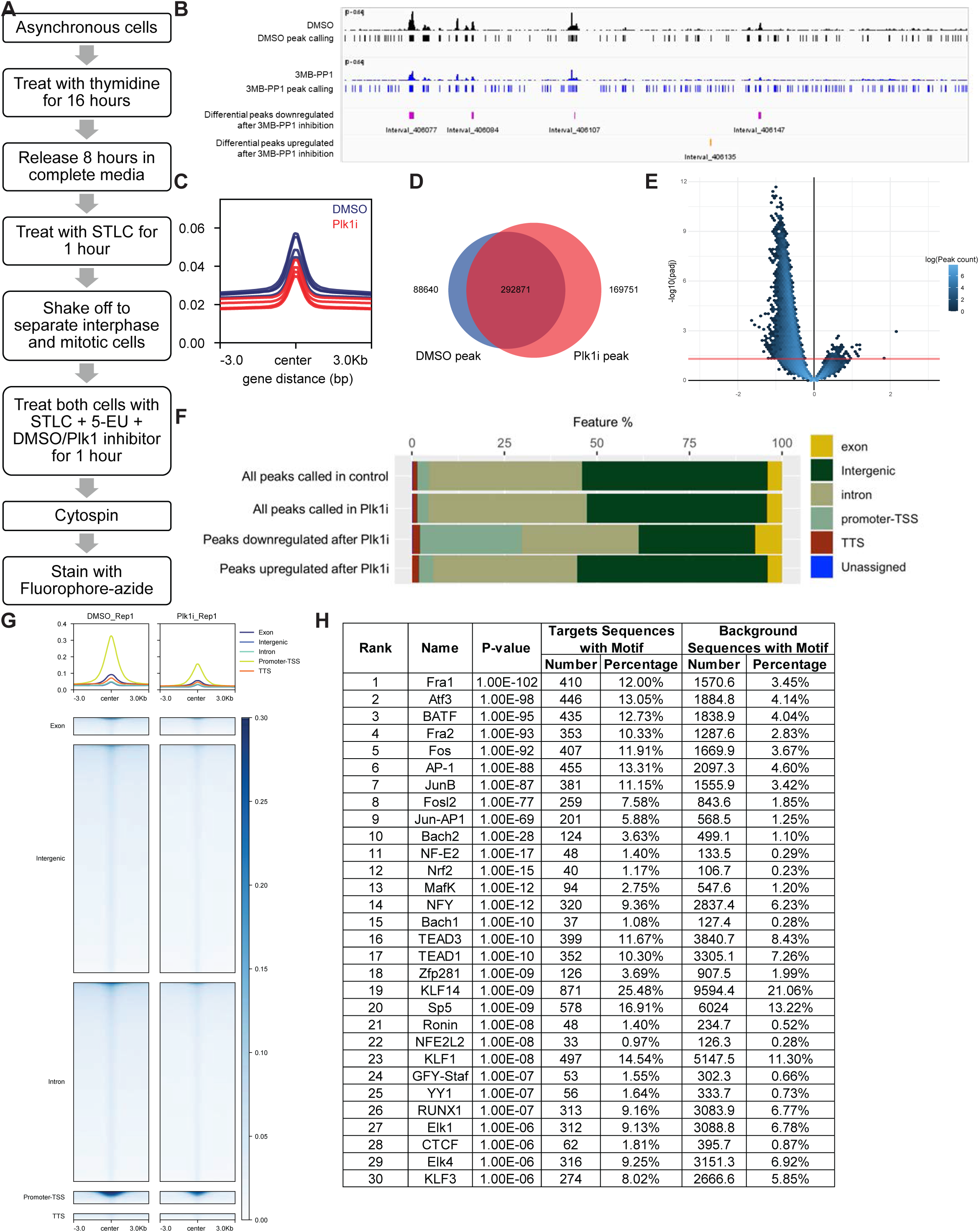
Plk1 Activity Regulates Chromosomal Accessibility in Mitosis. (A) Schematic of ATAC-seq in prometaphase arrested Plk1^AS^ cells after 2-hour DMSO or 10 µM 3MB-PP1 treatment. (B) IGV screenshot showed representative ATAC-seq peaks in control and Plk1 inhibited prometaphase Plk1^AS^ cells. Consensus, differential downregulated and upregulated peaks were shown. (C) Overlay of total mapped ATAC-seq peak profiles in control and Plk1 inhibited prometaphase Plk1^AS^ cells. (D) Venn diagram showed the merged peaks in control and Plk1 inhibited prometaphase Plk1^AS^ cells and the consensus peaks. (E) The volcano plot showed the differential expression of consensus peaks. Adjusted p-values was calculated using Benjamini-Hochberg method. Red line marked padj = 0.05. (F) Annotation profile of consensus and differential expressed peaks in control and Plk1 inhibited prometaphase Plk1^AS^ cells. (G) Representative ATAC-seq profiles of total peaks subsetted by gene annotation in control and Plk1 inhibited prometaphase Plk1^AS^ cells. (H) List of top 30 motif matched transcription factor records with enriched HOMER motif on differential peaks after Plk1 inhibition in prometaphase Plk1^AS^ cells. Adjusted p-value < 0.0001 (Benjamini-Hochberg method).

Compared to control cells, Plk1 inhibition at prometaphase resulted in an overall decrease in chromosomal accessibility characterized by reduced peak numbers and intensity, supporting our hypothesis (Fig. 4C, Fig. S4D). Mono-nucleosome sized fragments (180–247bp) were reduced in Plk1i group compared to control, suggesting reduced accessibility and increased condensation (Fig. S4A). Consensus peaks comprise 76.8% of total merged peaks mapped in the DMSO control group and 63.3% of the Plk1-inhibited group (Fig. 4D). Of 292,871 consensus peaks shared between both groups, 11,628 were significantly changed (fold change >2, p<0.05) after Plk1 inhibition, of which 99.50% are downregulated (Fig. 4E). This indicates that Plk1 activity is crucial for maintaining overall mitotic chromosomal accessibility. Further analysis revealed Plk1’s effects on the accessibility of specific chromatin regions. Annotating ATAC peaks into different functional categories revealed that peak distributions in Plk1-inhibited cells were similar to those in control cells, while significantly downregulated peaks showed an enrichment in promoter or transcription start site (promoter-TSS) regions (Fig. 4F). Subsetting peaks based on their genomic locations indicated a decrease in the promoter-TSS subgroup in both peak numbers and height (Fig. 4G). This suggests that Plk1 activity is crucial for maintaining the open chromatin status around promoter-TSS regions. Indeed, Homer Known Motif Enrichment analysis on peaks significantly reduced after Plk1 inhibition revealed an enrichment of binding motifs of key transcription factors and chromatin remodelers (Fig. 4H). Many transcription factors identified are in AP-1 superfamily (Fig. S4E). Therefore, the loss of Plk1 activity in prometaphase cells leads to the failure to retain open chromatin status at these genomic locations, preventing chromosomal interacting proteins from accessing the genome. Our findings indicate that Plk1 activity maintains the accessibility of certain promoter-TSS regions in mitotic chromosomes, thereby allowing transcription factors and machinery to initiate transcription. In conclusion, we propose a model whereby Plk1 activity restrains chromosome condensation during mitosis, maintaining chromosomal accessibility, particularly at promoter-TSS regions, thus facilitating transcription.

### Plk1 Activity Affects Phosphorylation of Condensin Proteins

The chromosome morphology changes observed after Plk1 inhibition resemble previously reported effects of condensin complex depletion (Elbatsh et al., 2019; Ono et al., 2003; Poser et al., 2020; Samejima et al., 2012). To further discern the molecular mechanism by which Plk1 influences the chromosomal landscape, we investigated the role of condensin complexes in Plk1-regulated chromosome activities. We utilized an inducible SMC4 knockdown cell line (HeLa SMC4-mAID-Halo, AAVS-OsTIR1(F74G) (Schneider et al., 2022), short for SMC4-AID), where endogenous SMC4 is rapidly degraded upon 5-PhIAA addition, as indicated by decreased Halo tag signal in immunoblots and time lapse imaging (Fig. 5A-B). As expected, Plk1 inhibition with BI-2536 reduced mitotic transcription in cells arrested at prometaphase (Fig. 5C-D). We noticed a population of cells with very low 5-EU incorporation in both group, likely due to the synchronization difficulties in this cell line.

**Figure 5.**
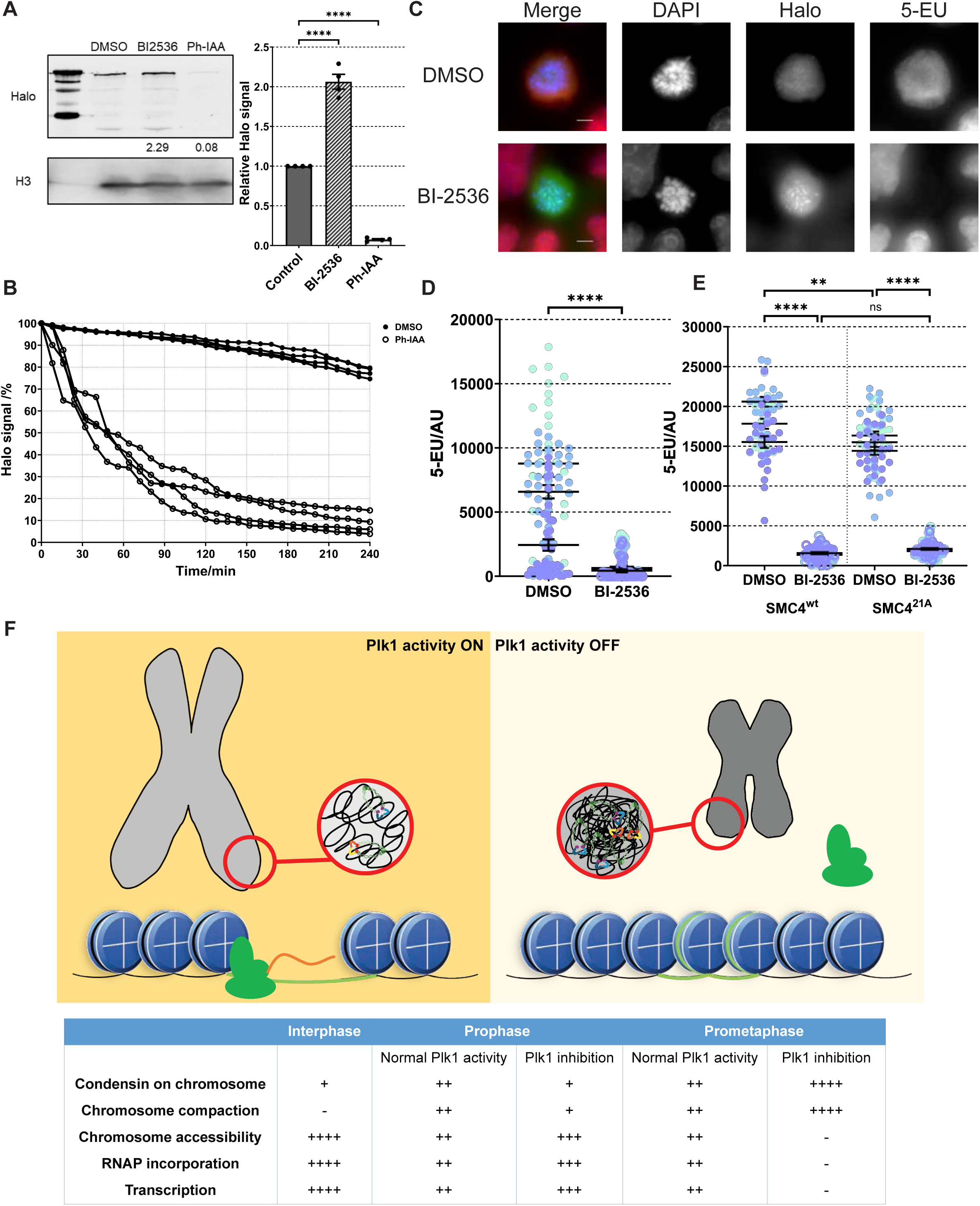
Plk1 potentially phosphorylates SMC4. (A) Western blot of chromosomal bound protein levels in mitotic HeLa SMC4-mAID-Halo, AAVS-OsTIR1 (SMC4-AID) cells. SMC4 level is assayed by HaloTag antibody. Histone H3 served as a loading control. Western blot band signals were quantified in the bar graph. (B) Time lapse of SMC4 degradation in SMC4-AID cells after addition of 5-PhIAA. SMC4 levels were assayed by HaloTag Oregon Green Ligand. (C) Representative images of SMC4-AID prometaphase cells after treatment treated with DMSO or 200 nM BI-2536 for 2 hours. (D) Quantification of chromosomal 5-EU fluorescent signal in SMC4-AID prometaphase cells showed in (C). (E) Quantification of chromosomal 5-EU fluorescent signal in prometaphase SMC4^WT^ and SMC4^21A^ cells treated with DMSO or 200 nM BI-2536 for 2 hours. (F) Model for Plk1 regulating mitotic chromosomal condensation, accessibility and transcription. During mitosis, after chromosome formation in prophase, Plk1 activity is essential for regulating the level of mitotic chromosome condensation. Plk1 phosphorylates substrates at chromosomal arms, potentially including the condensin component SMC4. This phosphorylation ensures the accessibility of chromatin regions within mitotic cells, thereby allowing transcription to occur during mitosis. Inhibition of Plk1 disrupts the equilibrium of condensin phosphorylation and chromosomal loading, leading to further chromosomal compaction. Consequently, chromosomal accessibility is reduced, preventing the transcription machinery from accessing and transcribing the mitotic genome, which results in impaired mitotic fidelity and likely affects post-mitotic events. For microscopic images, scale bar: 5 µm. For western quantification, each dot represents a biological replicate. For 5-EU quantification, each dot represents a cell, and different colors represent different biological replicates. 3 or more biological replicates were examined.

Previous research has shown that Plk1 phosphorylates condensin II-specific subunits CAP-D3, CAP-H2, and CAP-G2 (Abe et al., 2011; Kagami et al., 2017). Mutating Plk1 phosphorylation sites on CAP-D3 and CAP-H2 disrupts condensin II function and leads to improper mitotic segregation, confirming the critical role of Plk1 phosphorylation in regulating chromosomal assembly and ensuring mitotic fidelity (Abe et al., 2011; Kagami et al., 2017). However, the effect of Plk1 phosphorylation on condensin II components occurs in prophase, whereas the chromosomal over-condensation, transcription inhibition, and anaphase segregation defects we examined above were observed in later mitotic stages. Therefore, Plk1 regulation of known condensin II substrates cannot fully explain these phenotypes. Phosphoproteomic analyses, including work done by our group (Johnson et al., 2020; Santamaria et al., 2011), have identified condensin II-specific subunits as putative mitotic Plk1 substrates. Additionally, the SMC4 protein, a subunit shared between both condensin complexes (Takemoto et al., 2004), is identified in the screen as potentially phosphorylated by Plk1 in mitosis. If SMC4 is indeed a Plk1 substrate, Plk1 likely affects the mitotic functions of both condensin complexes through SMC4 phosphorylation.

To directly link Plk1 function with SMC4 phosphorylation, we conducted a rescue experiment in the SMC4-AID cell line (Fig. S5A). Briefly, we mutated 21 serine/threonine residues harboring potential Plk1 phosphorylation motifs in SMC4 into alanine to block any potential Plk1 phosphorylation effects. We then transfected mCherry-tagged wild-type SMC4 (SMC4^WT^) and the mutant (SMC4^21A^) into SMC4-AID cells where endogenous SMC4 can be degraded upon 5-PhIAA addition. After 5-PhIAA addition, we predicted that SMC4^21A^-transfected cells would exhibit lower basal levels of mitotic transcription compared to SMC4^WT^-transfected cells, indicating that Plk1 regulates mitotic transcription through SMC4 phosphorylation. We also predicted that mitotic transcription would not be further reduced by Plk1 inhibition in SMC4^21A^-transfected cells, indicating that SMC4 is the main target of Plk1 in regulating transcription. Both SMC4^WT^ and SMC4^21A^ cells remained viable after endogenous SMC4 degradation by 5-PhIAA addition. Transgene expression indicated by mCherry signal were observed in immunoblots and immunofluorescence (Fig. S5B-D). Compared to the parental cell line, both cells showed a slower growth, with SMC4^21A^ exhibiting a slight growth defect compared to SMC4^WT^ cells (Fig. S5E). When we treated both cell lines with STLC and 5-EU to label nascent transcription, prometaphase SMC4^21A^ cells showed a slight but statistically significant decrease in base-level 5-EU incorporation signal compared to SMC4^WT^ under the same conditions (Fig. 5E). This suggests that blocking Plk1 phosphorylation on SMC4 partially mimics the effect of Plk1 inhibition on mitotic transcription. Adding the Plk1 inhibitor BI-2536 reduced mitotic transcription levels in both cell lines to a uniform level, indicating that SMC4 is not the only substrate involved in Plk1-mediated mitotic transcription regulation (Fig. 5F). This aligns with previous findings of multiple Plk1 substrates within condensin complexes.

To summarize, we showed that SMC4 is a potential Plk1 substrate, and its phosphorylation by Plk1 restricts the degree of chromosomal condensation, allowing persistent transcription during human mitosis.

## Discussion

Contrary to the classical view suggesting that mitotic cells exhibit fully compacted chromosomes and are transcriptionally silent, it is now recognized that cells maintain a minimal level of transcription during mitosis. Substantial progress has been made in elucidating the biological functions of mitotic transcripts, which include both coding and non-coding RNAs, during and post-mitosis. However, the mechanisms sustaining transcription during mitosis remain poorly understood. Some studies have proposed that specific transcription factors, known as bookmarking factors, retain their association with mitotic chromatin to facilitate the post-mitotic transcription reactivation process. This regulation through transcription factor binding is regional and cannot fully account for chromosomal-level transcription.

Our study identifies Polo-like kinase 1 (Plk1) as a critical regulator of global mitotic transcription (Fig. 5F). As mitosis progresses, Plk1 activity restricts chromosome condensation, likely via condensin phosphorylation, to maintain chromosomal accessibility, thereby enabling transcription on the mitotic genome. In the case of Plk1 inhibition, condensin phosphorylation status is disrupted after prophase, which leads to over-condensation of chromosomes in later mitosis. This over-condensation hinders access to open chromatin regions, thereby reducing mitotic transcription. This discovery introduces a new layer of regulation for mitotic transcription, beyond the localization of transcription factors and RNA polymerases. Plk1 activity is essential for keeping chromosomal regions accessible, allowing the binding of transcription machinery, and thereby laying the foundation for downstream regulation of mitotic transcription in a more precise and cell-type-specific manner.

Previous studies revealed that transcription at early mitosis resulted from genes with engaged RNAP II to proceed transcription until the 3’-end of the gene (Liang et al., 2015). Proper chromosome segregation requires cohesin removal and RNAP II clearance from chromosomal arms during prophase (Perea-Resa et al., 2020). Plk1 inhibition at prophase, which prevents cohesin removal, results in the retention of elongating RNAP II on mitotic chromosome arms, corresponding to sustained transcription. However, by arresting cells in prometaphase prior to kinase inhibition and 5-EU labeling, we bypassed prophase to investigate transcription at a later mitotic stage and the regulatory effect of Plk1 activity. 5-EU staining on chromosome spreads indicated transcription in prometaphase cells, particularly on chromosomal arms (Fig. 1D-E, Fig. S1E-F), consistent with previous data (Palozola et al., 2017). Plk1 inhibition at prometaphase led to a marked reduction in the 5-EU signal, illustrating Plk1 activity’s necessity for normal mitotic transcription. These contrasting effects at different mitotic stages suggest the complexity of Plk1’s regulatory network.

Accumulated evidence suggests that mitotic transcription plays an essential role in ensuring mitotic fidelity, supporting centromeric functions, and facilitating post-mitotic transcription reactivation. α-amanitin treatment resulted in increased chromosome segregation errors in anaphase (Fig. 2A-C). Given that α-amanitin is a slow-acting drug, we were unable to determine whether transcription inhibition occurred before or after mitotic entry. Although some rapid-acting RNAP II inhibitors were employed in other studies and showed effects on mitotic progress (Bury et al., 2020; Chan et al., 2012; Perea-Resa et al., 2020), their effects are under debate due to potential off-target consequences (Y. Chen et al., 2021; Liu et al., 2015; Novais-Cruz et al., 2018). We utilize POLR2A-AID cells to deplete RNAP II component to suppress transcription, resulting in the similar chromosome segregation defects (Fig. 2D-E). Our research confirms that transcription is crucial for accurate chromosome segregation during anaphase. Notably, partial Plk1 inhibition results in a high incidence of chromosome missegregation, similar to the effect of transcription inhibition (Fig. 2A-E), underscoring Plk1’s critical role. Plk1 activity on chromosomes is essential for transcription and segregation regulation, as restoring Plk1 activity via expression of the Plk1 kinase domain fused with histone H2B partially rescues these phenotypes (Fig. 2G-H). Although we do not specifically assess the impact of Plk1 on centromeric transcription, previous findings from our lab demonstrated that a CENP-A-Plk1 kinase domain fusion protein failed to rescue the transcriptional and segregation phenotypes observed upon Plk1 inhibition (Lera et al., 2016, 2019). RNAP II inhibition causes overall transcription reduction, consistent with our observation of Plk1’s requirement on DNA (Fig. 2A-E). This indicates that transcriptional activity on chromosomal arms, rather than solely at centromeres, is critical for proper mitotic progression.

Although our experiments did not specifically analyze transcripts, prior studies have demonstrated that mitotically enriched genes are involved in extracellular structure and transcription processes, with some exhibiting prioritized transcription at mitotic exit (Palozola et al., 2017). Given the extensive chromosomal condensation and accessibility changes induced by Plk1 inhibition, we predict that the transcription of mitotically enriched genes is significantly impacted. Our results support the hypothesis that the loss of Plk1 activity impairs mitotic transcription, consequently inhibiting post-mitotic transcription reactivation. However, due to the necessity of Plk1 activity during mitotic exit and cytokinesis, we did not assess this effect directly. Future research should focus on achieving precise temporal control of Plk1 activity in late mitosis to further investigate this hypothesis.

Previous studies have shown that Plk1 regulates condensin II subunits CAP-D3 and CAP-H2 via phosphorylation in early prophase, which is critical for condensin II function (Abe et al., 2011; Kagami et al., 2017). Blocking Plk1 phosphorylation of CAP-D3 and CAP-H2 resulted in the retention of chromosomal condensation. On the other hand, we showed that in later mitotic stages, Plk1 inhibition led to significant over-condensation chromosomes, suggesting that Plk1 activity is required to restrict chromosome condensation level (Fig. 3). Together, results reinforce Plk1’s multifaceted role in regulating mitotic process and the requirement of delicate spatial-temporal regulation of kinase activity to ensure proper mitosis.

In addition to condensin II-specific components, we demonstrate that mutating potential Plk1 phosphorylation sites on SMC4 partially mimics Plk1 activity loss, and further Plk1 inhibition in SMC4 mutant and wild-type cells decreases mitotic transcription to the same baseline level (Fig. 5E). This confirms the involvement of other Plk1 substrates in regulating mitotic transcription. It is likely that Plk1 phosphorylates additional proteins involved in chromosome condensation, creating a balanced state of chromatin condensation that permits low but consistent mitotic transcriptional activity.

The underlying mechanisms regulating chromosomal accessibility remain poorly understood. Our findings demonstrate that Plk1 activity is pivotal in controlling chromosomal accessibility during mitosis, likely through interacting with condensin complexes. Using ATAC-seq, we observe that Plk1 inhibition reduces chromosomal accessibility not only at the global level, but also a significant downregulation at promoter-TSS regions (Fig. 4C-G). This indicates the existence of additional local regulatory mechanisms. It is possible that other Plk1 interactors contribute to the regulation of accessibility at specific genomic regions, or alternatively, that condensin proteins are recognized and regulated differentially across various genomic regions. Our research indicates that Plk1’s role in regulating chromosomal accessibility during mitosis involves a complex interplay between Plk1 and other chromatin-associated proteins.

Our findings provide the first evidence of the role of Plk1 activity in transcription regulation during mitosis. We demonstrate that Plk1 exerts the same regulatory effect on mitotic transcription in both non-transformed and transformed cell lines, indicating this mechanism is fundamental to cellular function. This discovery underscores Plk1’s crucial role in mitotic processes and highlights the importance of Plk1 in maintaining cellular homeostasis and genomic stability. Future research will focus on identifying specific substrates and interactors targeted by Plk1 to regulate chromosomal condensation, accessibility, and transcription during mitosis. These efforts will enhance our understanding of Plk1’s diverse functions and its potential as a therapeutic target in diseases characterized by dysregulated mitosis.

## Supporting information

Reply to reviewers_with original comments

## Acknowledgments

We also would like to thank Drs. William Sugden, Beth Weaver, Aussie Suzuki, Darcie Moore, Ramya Varadarajan, Roshan Norman for suggestions and feedback. We also thank Shermineh Bradford for helpful discussions. We thank Kazuhiro Maeshima for DLD1 POLR2A-AID cells and Maximilian Schneider for SMC4-AID cells. This work was supported by R01GM131068 and R01CA234904 to M.E.B., the UW Carbone Cancer Center (P30CA014520) and the Holden Comprehensive Cancer Center (P30CA086862)

## Methods

### Cell culture

All cell lines were maintained at 37°C and 5% CO_2_ in a humidified incubator and propagated in the following media supplemented with 10% fetal bovine serum (Avantor) and 100 units/ml penicillin–streptomycin (Gibco, Thermo Fisher): hTERT-immortalized human retinal pigment epithelial-1 (RPE1) cells and its derivative cells (Plk1^WT^, Plk1^AS^ and flag tagged fusion rescue cell lines), DMEM/F12 media (Thermo Fisher); HeLa, Cal-51, and Phoenix cells, DMEM/High Glucose (Cytiva HyClone), DLD1 tet-OsTIR1 mClover-mAID-POLR2A (POLR2A-AID), RPMI1640 (Sigma).

HeLa SMC4-mAID-Halo, AAVS-OsTIR1 (F74G) (SMC4-AID) was cultured in the same media as HeLa, with additional 1% GlutaMAX (Gibco), 0.5μg/mL Puromycin, and 6μg/mL Blasticidin S (Gibco, Thermo Fisher). Cells transfected with mCherry-tagged SMC4 were selected under 0.4mg/mL G418 (Gibco, Thermo Fisher). Final concentration of 1μM 5-PhIAA was added to maintain cells with endogenous SMC4 degraded.

Cells were passaged every 3-4 days at 80-90% confluency. All cell lines were tested for mycoplasma contamination with the MycoAlert Mycoplasma Detection Kit (Lonza).

For fixed imaging analysis, cells were plated on #1.5 glass coverslips (Fisherbrand) 3 days before use for experiments at 80-90% confluency. For time lapse imaging analysis, cells were plated on 24-well glass bottom plate with high performance #1.5 cover glass (Fisherbrand).

### Primary antibody and chemicals used

Plk1 (Santa Cruz, sc-17783, 1:500), mCherry (Takara Bio, 632496, 1:500), ACA (Immunovision, HCT-0100, 1:1000), FLAG (Sigma, F1804, 1:500), GFP (Thermo Fisher, A-11120, 1:500), Halo (Promega, G9281, 1:1000), H3 (Cell Signaling, 14269, 1:1000) 5-EU (Vector Laboratories, CCT-1261-25), AzDye 594 (Vector Laboratories, CCT-1295), Alexa Fluor 488 Azide (Thermo Fisher, A10266), BI-2536 (VWR, S1109), 3MB-PP1 (Fisher, 52-958), STLC (Santa Cruz, sc-202799), Thymidine (Millipore Sigma, 6060), GSK461364 (MedChemExpress, HY-50877), IAA (Thermo Fisher, AAA1055614), PhIAA (MedChemExpress, HY-134653), α-amanitin (Santa Cruz, sc-202440A)

### Cell line construction

Retroviral plasmid backbone pQCXIN was purchased from Clontech. pQCXIN-mCherry plasmid was generated in lab and used as the backbone. SMC4 wildtype and mutant cDNA was synthesized by Twist Bioscience in five fragments. Fragments and linearized backbone were assembled through Gibson assembly.

For stable retroviral transduction, constructs were co-transfected with a VSV-G envelope plasmid into Phoenix cells. Fresh medium was applied 24 hours post-transfection. Another 24 hours later, virus-containing media were centrifuged and filtered through a 0.45μm membrane to remove cell debris, then diluted 1:1 with complete medium containing 10μg/mL Polybrene (Millipore). Target cells were infected at 40-60% confluency for 24 hours and then selected with 0.4mg/mL G418 for 10-14 days.

### Nascent transcript labeling

1mM 5-ethyl uridine (5-EU, Click Chemistry Tools, Vector Laboratories) were added to cells along with STLC (Santa Cruz) and drugs during 2-hour treatment at 37°C. Cell were plated or spun (see next section) onto coverslips before 5-EU click reaction.

Coverslips were fixed in 4% paraformaldehyde (PFA) in Phosphate-buffered saline (PBS) buffer for 10 min at 37°C, washed three times in PBS, permeabilized in PBS+0.5% Triton-X for 10 min at 37°C, washed three times in 0.1% Triton X-100 in PBS (PBST). Coverslips were then incubated in click reaction buffer (final concentration: 43 mM Tris, 129 mM NaCl, 1 mM CuSO_4_, 5.6mM sodium ascorbate, 9.6μM azide (Alexa Fluor 488 Azide, Thermo Scientific or AZDye 594 Azide, Click Chemistry Tools, Vector Laboratories)) in the dark at room temperature for 1 hour before washed in 0.5mM EDTA, 2mM NaN3 in PBS. Coverslips were washed three times with PBST, counterstained with DAPI (Sigma-Aldrich), mounted on glass slides with Prolong Diamond antifade medium (Molecular Probes), and allowed to cure for 48 hours.

### Chromosome spread

Cells were synchronized through a 16-hour STLC treatment before mitotic shakeoff and were subjected to their respective drug treatment for 2 hours. Cells were collected by centrifugation and resuspended in a 0.075M KCl hypotonic solution for 30 minutes at 37°C. For 5-EU labeled mitotic cells, cells were spun onto coverslips at 800 rpm for 5 minutes at medium acceleration (Cytospin 4, Epredia, Thermo Fisher). Coverslips were then incubated in KCM buffer for 10 minutes at room temperature before fixation. For chromosome condensation analysis, swollen cells were resuspended in fresh fixative (methanol: glacial acetic acid, 3:1 (v/v)). After three rounds of centrifuging and resuspension in fixative, cells were dropped onto cold tilted slides for spread. Slides were stained with DAPI, mounted, cured and subject to imaging.

### Immunofluorescence

Coverslips were fixed in 4% paraformaldehyde (PFA) in phosphate-buffered saline (PBS) buffer for 10 minutes at 37°C, washed three times in PBS, permeabilized in PBS+0.5% Triton-X for 10 minutes at 37°C, washed three times in 0.1% Triton X-100 in PBS (PBST). Coverslips were then incubated in click reaction buffer (final concentration: 43 mM Tris, 129 mM NaCl, 1 mM CuSO4, 9.6μM fluorophore azide, 5.6mM sodium ascorbate) in the dark at room temperature for 1 hour before being washed in 0.5mM EDTA and 2mM NaN3 in PBS.

Coverslips were washed three times with PBST, counterstained with DAPI (Sigma-Aldrich), mounted on glass slides with ProLong Diamond antifade medium (Molecular Probes), and allowed to cure for 48 hours.

### ATAC-seq

We processed ATAC-seq samples using a protocol modified from Buenrostro et al., 2013. As shown in Fig. 4A, prometaphase RPE1 Plk1^AS^ cells were synchronized in 5 µM STLC treatment for 16 hours and got isolated by mitotic shakeoff. Cells were treated with DMSO or 10 µM 3MB-PP1 in STLC containing media for 2 hours. Before collection, mitotic index of each sample was measured (>95%). Four biological replicates were achieved. For each condition, 50,000 cells were resuspended in 50 uL transposition reaction mix containing 25 uL Tagment DNA buffer (Illumina), 16.5 uL 1x PBS, 0.5 uL 10% Tween-20, 0.5 uL 1% Digitonin, 2.5 uL Tn5 Transposase (Illumina), and 5 uL nuclease-free water. The reaction mixture was incubated at 37°C for 30 min in a Thermomixer set at 1000 rpm. DNA was isolated using a QIAquick PCR Purification kit (Qiagen) according to manufacturer’s protocol, and DNA was eluted in 10uL of the supplied elution buffer. The entirety of the purified DNA was added to a PCR reaction containing 10 uL nuclease-free water, 2.5 uL each of 25uM NextEra i5 and i7 barcoded primers (Active Motif), and 25 uL NEBNext high-Fidelity 2X PCR Master Mix (New England Biolabs).

Samples were amplified on the following program: 72°C 5 min, 90°C 30 sec, (98°C 10 sec, 63°C 30 sec, 63°C 1 min) x 5 cycles. 5 uL of the PCR reaction was removed and added to 3.85 uL nuclease-free water, 0.5 uL each of i5 and i7 barcoded primers, 0.15 uL 100x SYBR Safe DNA stain (Invitrogen), and 5 uL NEBNext High-Fidelity 2x PCR Master Mix. qPCR was performed using the following program: 98°C 30 sec, (98°C 10 sec, 63°C 30 sec, 72°C 1min) x 20 cycles. Following qPCR, the R value vs cycle number for each sample was plotted to calculate the number of cycles needed to reach 1/3 of the maximum R, and that number was used to continue PCR amplification of the libraries. This step is to stop amplification prior to saturation to avoid variation among samples due to PCR bias. After final PCR amplification, the samples were size-selected using SPRIselect beads (Beckman Coulter) by doing a 0.5x standard bead cleanup followed by a 1.8x standard bead cleanup. Samples were eluted from the beads using 20uL of nuclease-free water. Samples were analyzed on a TapeStation 4200 instrument using high-sensitivity D5000 ScreenTapes and reagents (Agilent) and quantified using high-sensitivity dsDNA reagents on a Qubit 4 fluorometer (Invitrogen). Samples were pooled equimolarly and sequenced to a target depth of 50 million paired-end reads on a NextSeq 2000 instrument (Illumina).

### ATAC-seq data analysis

Analysis of the sequencing data was performed using Nextflow’s ATACseq pipeline (DI Tommaso et al., 2017), which uses MACS2 (Zhang et al., 2008) for peak calling. The peaks from the DMSO and 3MB-PP1 treated samples were merged to form consensus peaks.

Differential ATAC signal was quantified over these consensus peaks using DEseq2 (Love et al., 2014). Profile plots were generated using DeepTools computeMatrix (Ramírez et al., 2016). Peak annotation was performed to identify what types of genomic regions were represented in the ATAC peaks (Benner et al., 2017) and was used to subset peaks in profile plots in Fig. 4F and Fig. S4D. Peak distribution is analyzed using Picard as part of the MultiQC report in the nextflow pipeline. Peak set similarity was plotted using the Intervene pairwise function to plot the jaccard similarity in Fig. S4B (Khan & Mathelier, 2017).

The cutoffs for HOMER analysis were determined by testing motif discovery on a variety of cutoffs by number of peaks with fold enrichment > 2. In the DMSO and 3MB-PP1 treated peaks, enriched motifs follow a similar pattern up through ∼4000 peaks and then by ∼8000 are dominated by background noise. There are only 59 significant differential peaks that are up in 3MB, so we have no reason to try varying the number of peaks. There are only ∼12,000 significant differential peaks that are downregulated in 3MB-PP1 treated group. The motif patterns stay consistent throughout iterations on the number of peaks included. Therefore, 4000 peaks with the highest signal were selected from each of the DMSO, 3MB-PP1 treated, and differential peaks to perform Homer Known Motif Enrichment analysis (Benner et al., 2017). String plot, Gene ontology enrichment and protein interaction was made using STRING (v12.0, Szklarczyk et al., 2023).

The detailed ATAC-seq data is uploaded on GEO database (GSE282486) and the code used can be found in Github: https://github.com/jessicalanglab/plk1_atac_analysis.

### Subcellular fractionation

Cells were synchronized through a 16-hour STLC treatment before mitotic shake-off and were subjected to their respective drug treatment for 2 hours before collection. Pelleted cells were lysed in HB2 buffer (50mM HEPES, pH 7.5, 0.5% NP-40, 10% glycerol, 100 mM NaCl, 10 mM Na pyrophosphate, 5 mM β-glycerophosphate, 50 mM NaF, 0.3 mM Na3VO4, 1 mM DTT, 1 mM PMSF, and 1x complete protease inhibitor cocktail (EDTA-free)) on ice, and lysate was centrifuged at 12000xg for 15 minutes at 4°C. Cytosolic proteins were concentrated in the supernatant, and nuclear proteins were enriched in the pellets. 4x Protein Sample Buffer (62.5mM Tris, 4% (w/v) SDS, 20% (v/v) glycerol, and 0.01% (w/v) bromophenol blue) was added and boiled for 5 minutes.

### Western blot

Proteins in the sample were separated by SDS–PAGE, transferred onto a 0.2 μm nitrocellulose membrane, and blocked for 1 hour in blocking buffer (5% milk and Tris-buffered saline pH 7.4 with 0.1% Tween-20 (TBST)). To simultaneously probe for the protein of interest and the loading marker, the membrane was divided in two after transfer and incubated in separate antibody solutions. Membranes were incubated with gentle agitation overnight at 4°C with primary antibodies diluted in blocking buffer, washed with TBST, and incubated for 1 hour at room temperature in IRDye secondary antibodies (LI-COR) diluted in blocking buffer.

Membranes were washed with TBST and rinsed once in TBS before being imaged in a LI-COR Odyssey Imager. Images were quantified in Image Studio (LI-COR).

### Growth assay

Cell growth was determined using CellTiter 96® Non-Radioactive Cell Proliferation Assay (MTT) (Promega). Briefly, cells plated in 96-well plates with corresponding culture conditions were grown for 5 days before collection. Leave 100 µL media in the well, add 15 µL Dye Solution. After a 2-hour incubation at 37°C, add 100 µL Solubilization Solution/Stop Mix to each well. Measure A570 and normalize to no cell control.

### Imaging and analysis

Image acquisition was performed on a Nikon Eclipse Ti inverted microscope equipped with a motorized stage, LED epifluorescence light source (Spectra X), 40x air objective or 100x oil objective, and ORCA Flash4.0 V2+ digital sCMOS camera (Hamamatsu).

For regular slides, prometaphase cells were determined by DAPI channel morphology under the microscope, and multiple z-stack images for each channel were taken. For spreads, evenly spread cells were imaged.

To quantify chromosomal fluorescence: after background normalization, chromosomal area was determined using a uniform threshold for DAPI signal across the same dataset.

Chromosomal fluorescent signal (5-EU or immunostaining) overlapping with chromosome area volume was measured. More than 20 cells were analyzed for each replicate, and at least 3 biological replicates were achieved.

To measure chromosome area: after background normalization, chromosomal area was determined using a uniform threshold for DAPI signal across the same dataset. More than 20 cells were analyzed for each replicate, and at least 3 biological replicates were achieved.

To measure mitotic errors: immunofluorescence slides were probed for Plk1 and ACA. Anaphase cells were determined by Plk1 localization under the microscope. DAPI and ACA were used to evaluate segregation. More than 50 cells were analyzed for each replicate, and at least 3 biological replicates were achieved.

To measure the Halo tag degradation: cells were plated and pretreated (if needed) in 24-well glass bottom plates. Before adding the drug, multiple xy points were determined. Images were collected every 5-10 minutes for 4 hours. Environmental control was maintained by a humidified, stage-top chamber (Tokei Hit) set to 37°C and 5% CO2. For analysis, nuclear intensity was measured in each time frame.

### Statistical analysis

Data analysis was performed using Prism 8 (GraphPad). Statistical significance was determined using a two-tailed, unpaired t-test with Welch’s correction when comparing a single cell line with a single drug treatment, or one-way ANOVA with Tukey or Dunnett multiple comparisons test when comparing multiple treatments, or two-way ANOVA with Tukey or Dunnett multiple comparisons test when comparing both different cell lines and different treatments. *, P < 0.05; **, P < 0.01; ***, P < 0.001; ns, not significant.

---

**Figure S1.**
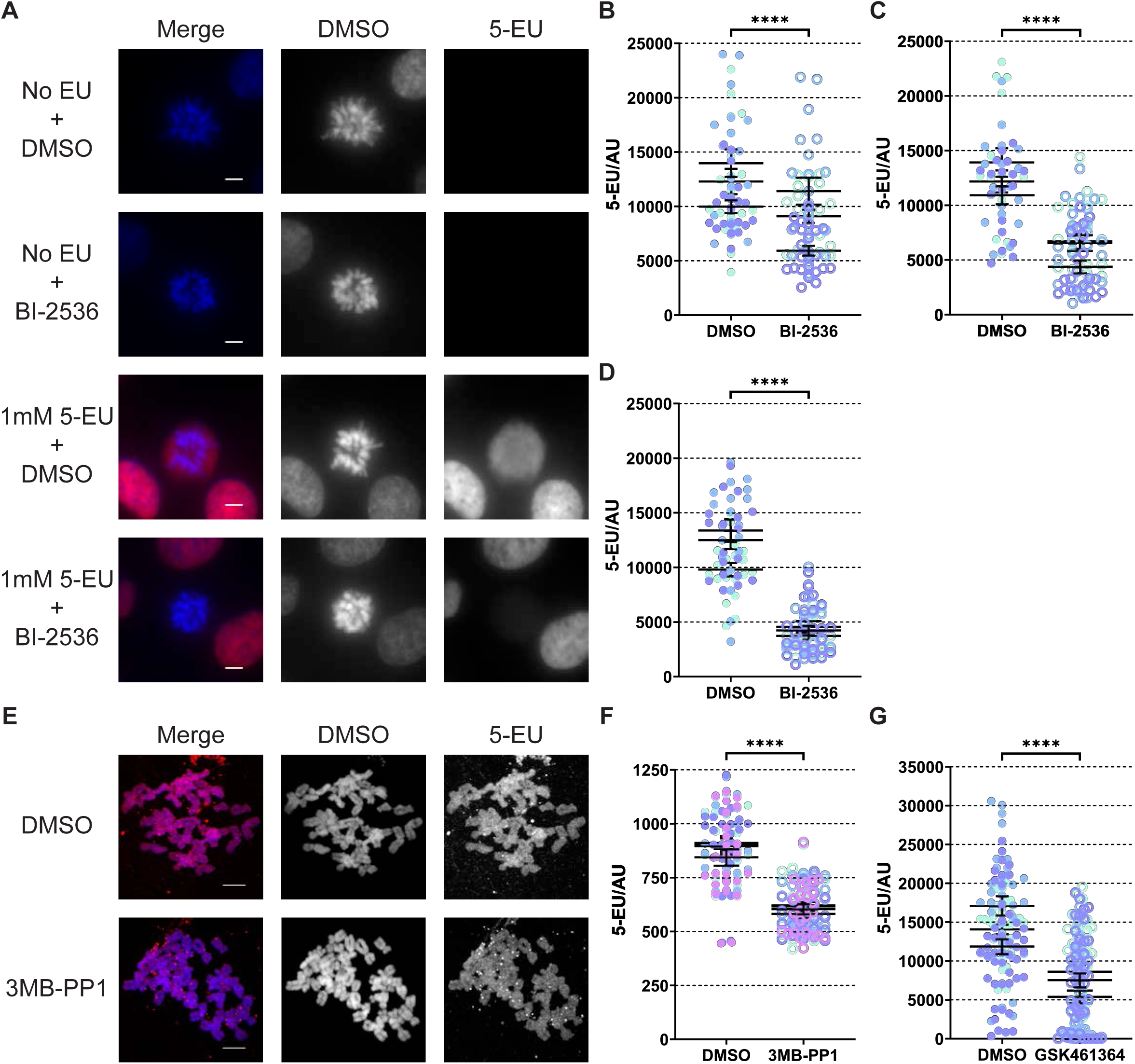
Effect of Plk1 activity on mitosis transcription in multiple cell lines. (A) Representative images of 5-EU staining corresponding to nascent transcription in prometaphase RPE1 cells with indicated drug treatment. (B) Quantification of chromosomal 5-EU fluorescent signal in prometaphase MDA-MB-231 cells treated with DMSO or 200 nM BI-2536 for 2 hours. (C) Quantification of chromosomal 5-EU fluorescent signal in prometaphase MDA-MB-453 cells treated with DMSO or 200 nM BI-2536 for 2 hours. (D) Quantification of chromosomal 5-EU fluorescent signal in prometaphase MDA-MB-468 cells treated with DMSO or 200 nM BI-2536 for 2 hours. (E) Representative images of Plk1^AS^ prometaphase cell spreads. Cells were synchronized in STLC and collected by shakeoff, followed by 2 hours DMSO or final concentration of 10 µM 3MB-PP1 in 1 mM 5-EU and 5 µM STLC containing media. (F) Quantification of chromosomal 5-EU fluorescent signal in Plk1^AS^ prometaphase cell spreads showed in (E). (G) Quantification of chromosomal 5-EU fluorescent signal in prometaphase RPE1 cells treated with DMSO or 1 µM Plk1 inhibitor GSK461364 for 2 hours. For microscopic images, scale bar: 5 µm. For 5-EU quantification, each dot represents a cell, and different colors represent different biological replicates. 3 or more biological replicates were examined.

**Figure S2.**
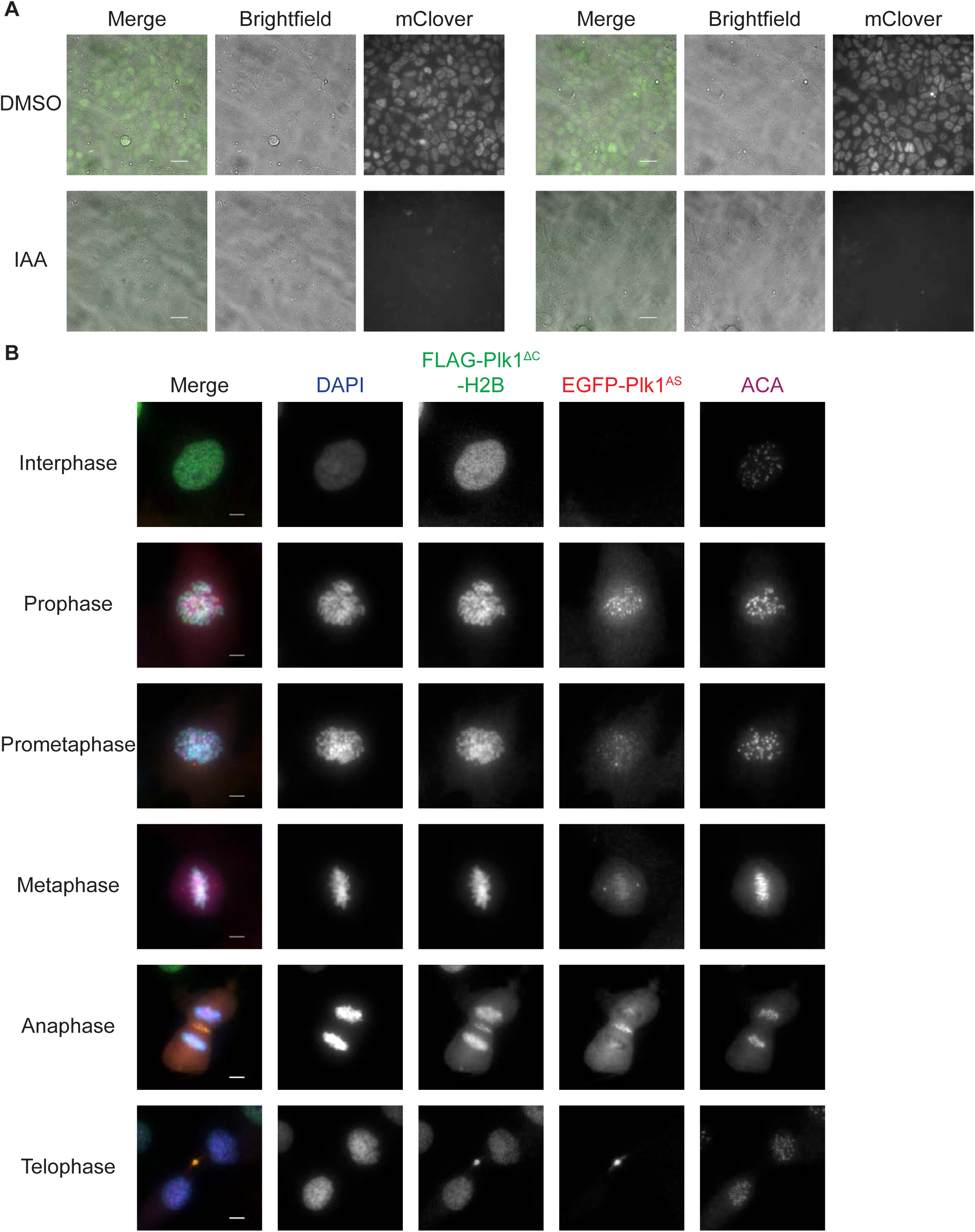
The effect of mitotic transcription and Plk1 activity on mitotic fidelity. (A) Representative images of DLD1 tet-OsTIR1 mClover-mAID-POLR2A cells pretreated for 23-hour 1 μg/mL doxycycline, and incubated in DMSO, 0.5 mM IAA for 3 hours. mClover signal suggested the POLR2A protein. Scale bar: 25 µm. (B) Representative images of RPE1 Plk1^-/-^ EGFP-Plk1^AS^ FLAG-Plk1^ΔC^-H2B (FLAG-Plk1^ΔC^-H2B) at each cell cycle stage. The FLAG signal suggested the chromosomal localization of Plk1 kinase domain throughout cell cycle. Scale bar: 5 µm.

**Figure S3.**
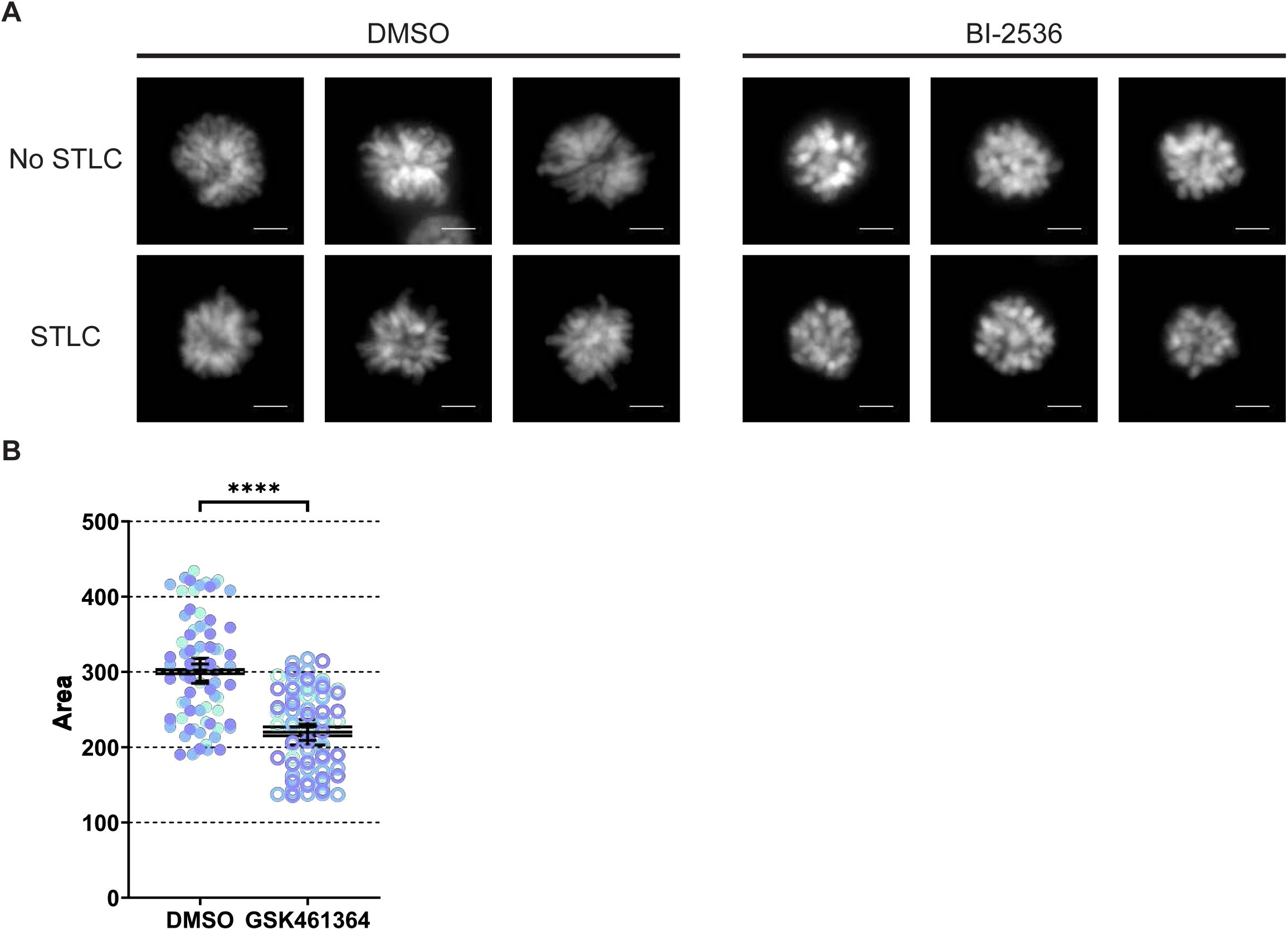
The effect of Plk1 activity on chromosome condensation. (A) Representative images of chromosomal arrangement in RPE1 cells treated with indicated drug treatment for 2 hours. (B) Quantification of chromosomal area in chromosomal spreads in RPE1 prometaphase cells treated with DMSO or 1 µM GSK461364 for 2 hours. For microscopic images, scale bar: 5 µm. For chromosomal condensation quantification, each dot represents a cell, and different colors represent different biological replicates. 3 or more biological replicates were examined.

**Figure S4.**
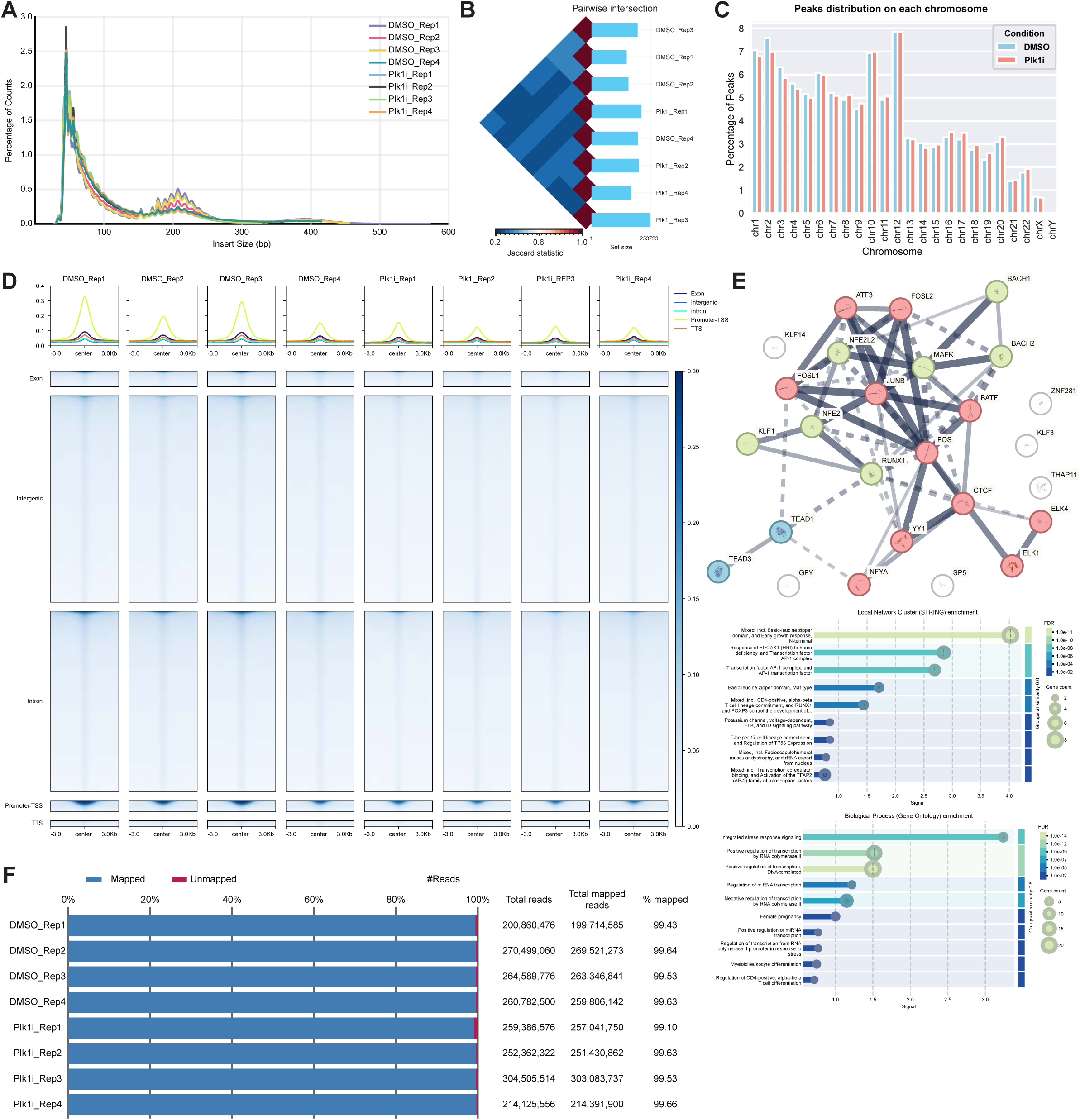
ATACseq analysis on mitotic cells affected by Plk1 inhibition. (A) Fragment length distribution of ATAC-seq reads for control and Plk1 inhibited prometaphase Plk1^AS^ cells. (B) Intervene diagram showed the ov’aerlapped peaks between replicates in each treatment group. (C) Distribution of consensus peak of control and Plk1 inhibited cells at each chromosome. (D) ATAC-seq profiles of total peaks in all replicates of control and Plk1 inhibited prometaphase Plk1AS cells. Peaks were subsetted by gene annotation. (E) Enriched transcription factors in HOMER analysis on differential downregulated peaks after Plk1 inhibition were plotted in a string plot. STRING network indicates both functional and physical protein associations. Network edges mean confidence (> 0.400). Line thickness indicates the strength of data support (from text mining, experiments, databases, co-expression, neighborhood, gene fusion, and co-occurrence). Clustering using k-means (find a defined number of clusters based on their centroids). 3 clusters were generated (Cluster 1, red, 11; Cluster 2, green, 7; Cluster 3, blue, 2). Edges between clusters were indicated by dotted line. (F) Summary of the ATAC-seq library.

**Figure S5.**
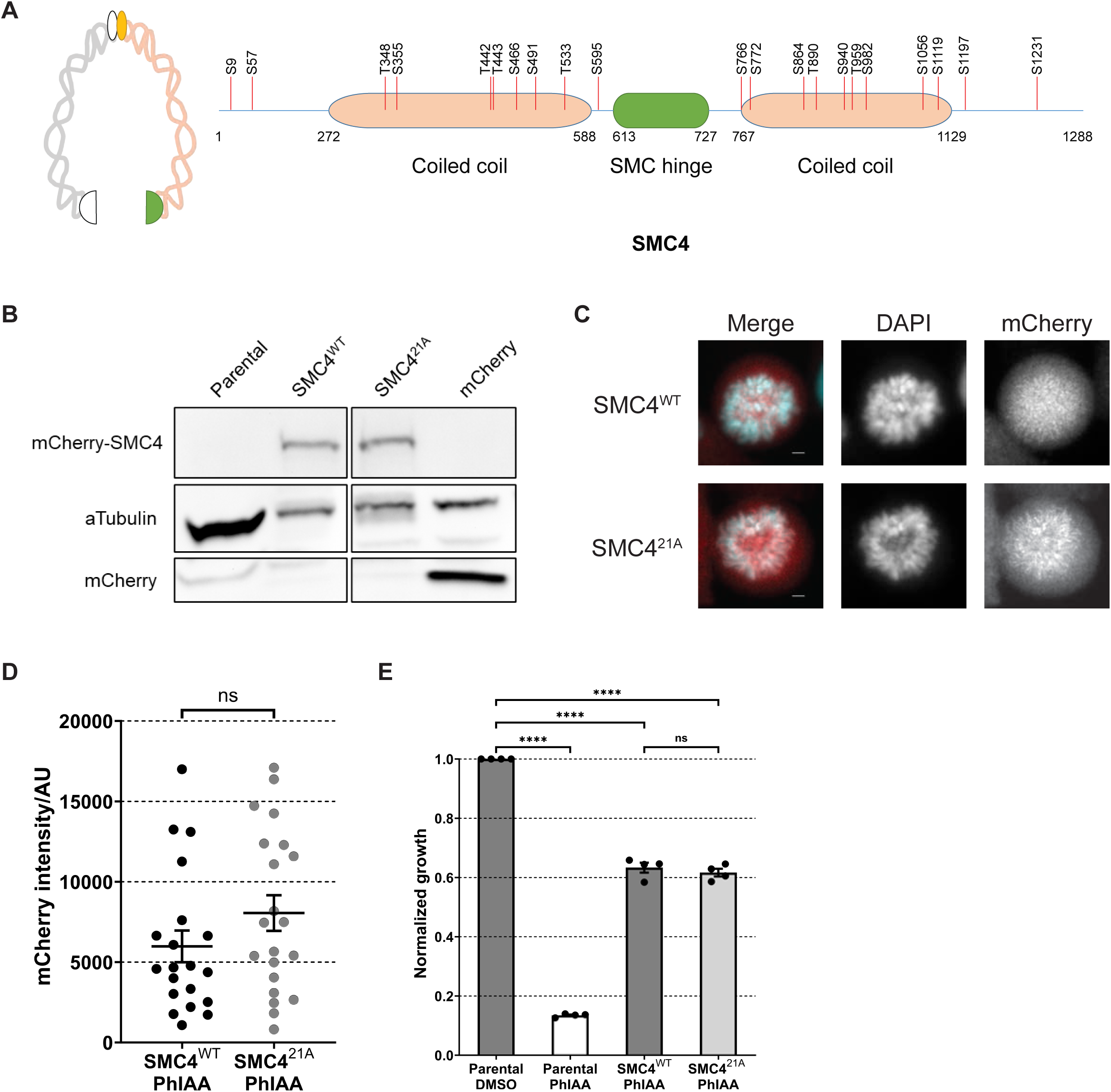
Validation of SMC4^WT^ and SMC4^21A^ cells. (A) Illustration of SMC4 protein domain structure and the design of the 21 sites mutated in Plk1 phosphor-blocking SMC4 mutant (SMC4^21A^). Mutated 21 residues: S9, S57, T348, S355, T442, T443, S466, S491, T533, S595, S766, S772, S864, T890, S940, T959, S982, S1056, S1119, T1197, S1231. (B) Western blot of total protein levels in SMC4^WT^ and SMC4^21A^ cells. (C) Representative images of prometaphase SMC4^WT^ and SMC4^21A^ cells. Scale bar: 5 µm. (D) Quantification of chromosomal mCherry signal in prometaphase cells showed in (B). Cells were incubated in regular media supplemented with 1µM PhIAA. No synchronization is done. Each dot represents a cell, 1 replicate was examined. (E) Growth assay of SMC4^WT^ and SMC4^21A^ lines. Each dot represents a biological replicate, 3 or more biological replicates were examined.

