## Supplementary material for "Polo-like kinase 1 maintains transcription and chromosomal accessibility during mitosis": Reply to reviewers_with original comments

**Reviewer 1:**

Major points:

1. To show a lack of transcription in mitotically arrested Plk1-inhibited cells, the authors perform a 2 hour treatment with STLC together with a pulse of 5EU (Fig 1A). As the treated cells would not immediately enter and arrest in mitosis upon drug addition, this treatment will result in labeling not only of mitotically transcribed RNA, but also transcripts synthesized in G2 cells before they arrest in mitosis due to STLC treatment. As Plk1 is also active G2, it is important for the authors to address this possibility. This issue is avoided in other experiments that include the addition of a shake-off step prior to drug treatment (eg. Fig 3B). This can be addressed by adding a shake-off step (potentially with a spin to get the cells back onto the plate for imaging?) or at least by acknowledging the possible contribution of G2-synthesized transcripts to the EU signal in the text. There is also a similar issue in Fig 4C-D / lines 280-281

*We agree on the importance of considering the effect of G2 transcripts in the 5-EU experiment, as discussed in the text. Specifically, we tested the effect of Plk1 activity on nascent transcription in mitotic and G2 phases in Figures 1C-E.*

2. The title of section 2 (line 172) implies that the authors demonstrate that chromosomal Plk1 activity is on-pathway with mitotic transcription to maintain mitotic fidelity. This is misleading since the experiments in this section show a correlation, but not a direct relationship (and in fact many perturbations alter mitotic fidelity). In addition, the authors never test the combined effect of Plk1 inhibition and transcriptional inhibition. We suggest the authors perform experiments with combined transcriptional inhibitor and Plk1 inhibitor treatments (perhaps titrating both treatments to assess the dose response), which would better justify the section title "Transcription maintains mitotic fidelity with a requirement for chromosomal Plk1 activity". Such an experiment would also strengthen the claim in lines 184-185.

*We acknowledge that our experiments only demonstrate a correlation between transcription and Plk1’s effect on mitotic fidelity. As suggested, we attempted double inhibitor treatment but under these conditions, the mitotic checkpoint was satisfied and cells didn’t enter anaphase. Consequently, we revised the section title to temper the statement.*

3. In Figure 3C-D, the authors show a decrease in chromosome area upon Plk1 inhibition. We think this experiment would benefit from having a positive control - another perturbation that is known to lead to overly condensed chromosomes. This would provide a sense of scale of the effect size observed in BI-2536-treated cells.

*Given this suggestion, we reviewed the literature for possible controls which would recapitulate the condensed phenotype and were unable to identify a suitable positive control. However, we demonstrated that Plk1 inhibition alone caused over-condensation.*

4. The speculation that Plk1 allows mitotic transcription by preventing over-condensation of chromosomes would be stronger if the authors showed that their Plk1dC-H2B construct can rescue the chromosome density phenotype. The experiments presented in this study do not definitively show this mechanistic pathway, so we urge the authors to be more reserved in their wording. (lines 240-241)

*We revised our wording to temper the statement.*

5. The authors claim that Plk1-inhibited samples showed fewer ATAC-seq peaks and reduced peak intensity based on Fig 3J. Our understanding of this figure is that, while it shows lower peak intensity, it does not demonstrate a difference in number of peaks, so authors should cite another figure or statistic for this claim. Additionally, given the reported statistics (only ~1/4 of the consensus peaks downregulated) the claim that Plk1 activity is crucial for maintaining overall mitotic chromosomal accessibility is too strong.

*We added a Venn diagram (Figure 4D) to illustrate the overlap of peaks between the control and Plk1-inhibited groups. more peaks were called in PLK1i group with their overall peak intensity decreased. Of 292,871 consensus peaks shared between both groups, 11,628 were significantly changed (fold change >2, p<0.05) after Plk1 inhibition, of which 99.50% are downregulated. Additionally, the peak profile (Figure 4C) and volcano plot (Figure 4E) highlight the overall downregulation after Plk1 inhibition.*

6. The role of Plk1-mediated phosphorylation of SMC4 in regulating mitotic transcription is overstated in the text. For example, there is no data showing that the phosphorylation of SMC4 by Plk1 restricts the degree of chromosomal condensation. The authors should perform chromosome area measurements in chromosome spreads, as in Fig 3C-G, or remove this claim (lines 305-306). In addition, the claim that SMC4 phosphorylation by Plk1 allows persistent transcription during human mitosis is too strongly worded given the demonstrated effect size in Fig 4F. Similarly, we found the role of condensin phosphorylation in discussion to be overstated given the limitations of this study (lines 320-322, line 347)

*We revised our wording to temper the statement.*

Minor points:

1. The introduction is quite long, and would benefit from some streamlining to focus the reader on the most critical points. In addition, many of points cited in the introduction - such as the role of transcription in CENP-A deposition) remain controversial, and it would be helpful to use slightly more cautious wording in describing this prior work and the potential roles of mitotic transcription. There are also serious prior issues related to off-target effects of many inhibitors that were used in prior studies (for example, see PMID: 30080136).

*We shortened part of the introduction.*

2. The article overall would benefit from some proofreading.

*We conducted a thorough proofreading of the manuscript.*

3. Fig 1C: the 6th box says both interphase and mitotic cells were treated with STLC. This is either a typo, or confusing experimental design. To arrest cells in G2, the authors could use the CDK1 inhibitor RO3306.

*To ensure consistency, G2 cells were treated with the same STLC-containing media as mitotic cells. However, only G2/interphase cells were used at later imaging and analysis steps.*

4. Line 162: missing reference to Fig 1H

*This issue has been corrected.*

5. Line 194: The naming of the H2B-tagged Plk1 mutant is inconsistent between text and figure.

*This issue has been corrected.*

6. Fig 3I-M: it would be interesting (although not necessary) to have some analysis of what genes undergo this Plk1-dependent regulation, not just which transcription factors may be affected.

*Thank you for the suggestion.*

7. Line 293: it is not appropriate to claim that Plk1 is the kinase regulating mitotic transcription through SMC4 phosphorylation just on the basis of this experiment. The 21A mutant may be disrupting other kinase target sites, not just Plk1. We suggest the authors rephrase this to "..indicating that mitotic transcription is regulated through SMC4 phosphorylation". Similarly, in line 300, there is insufficient evidence that Plk1 is the relevant kinase. This can be claimed in lines 301-302 when referencing the BI-2536 experiment, but not in line 300.

*We revised our wording to temper the statement.*

8. Fig 4D: there is a population of very low EU cells in the DMSO condition that was not present in other experiments. It would be helpful to comment on this in the text.

*This information has been mentioned in the text.*

9. Throughout the Figures, some plots have EU as y-axis label, but some have EU/AU. This should either be explained, or made consistent.

*This issue has been corrected.*

10. Supp Fig 2 lacks color legend.

*This issue has been corrected.*

11. Supp Fig 3A: please report the stain used.

*We included this information in the materials section.*

**Reviewer 2:**

Major points:

1. Regarding the title of the 3rd results section: "Plk1 Regulates Mitotic Transcription Through Chromosome Condensation". And also the last sentence: "They (these findings) also provide a mechanistic explanation for how Plk1 regulates mitotic transcription."

There is no evidence that the defect in chromosome condensation is a cause of the defect in transcription. In principle, to test it, one would have to somehow restore normal condensation in Plk1-inhibited cells and assay if mitotic transcription is rescued or not. This may not be possible experimentally in the current situation. If the authors eventually identify a key effector of chromosome condensation that is regulated by Plk1 in this process, the expression of a form of this protein with phosphomimetic mutations at Plk1 sites could be used. I don't think SMC4 is this factor (see point 3 below). Otherwise, it remains possible that the transcription defect observed after Plk1 inhibition is independent from its effect on chromosome condensation.

*We revised the title of the section and tempered the statement.*

2. Regarding the title "Plk1 Phosphorylates Condensin Proteins", it is not well chosen for this results section. Although its is true that Plk1 has been shown to phosphorylate some condensin subunits in previous publications, the current manuscript does not show any result in this regard.

*We revised the title of the section.*

3. The previous studies cited have reported a few phosphorylation sites in SMC4 that are dependent on Plk1 activity. However, mutating the 21 consensus sites in SMC4 for Plk1 is a questionable approach. Plk1 phosphorylation sites do not strictly match the proposed consensus motif, nor are the sites that match this motif in a protein always phosphorylated by Plk1. Importantly, the decrease in mitotic transcription observed when SMC421A is expressed compared to when SMC4WT is expressed in replacement of endogenous SMC4 is very small (a 10% to 15% decrease in Fig 4F). By contrast, when Plk1 is inhibited, the loss of mitotic transcription is almost complete (around 90%). Therefore, SM4C does not appear to be a major effector of Plk1 in the proposed mechanism. Moreover, the small effect observed could be due to a perturbation of condensin function due to the mutation of 21 sites that is unrelated to a loss of phosphorylation by Plk1. For example, SMC4 could be misfolded or destabilized by the mutations. No Western blot is shown to compare le levels of SMC421A and SMC4WT expressed.

*We included a western blot (Fig. S5A) to show mCherry-SMC4^WT^ and SMC4^21A^ expression.*

4. The Results section ends with this conclusion: "To summarize, SMC4 is a potential Plk1 substrate, and its phosphorylation by Plk1 restricts the degree of chromosomal condensation, allowing persistent transcription during human mitosis."

This conclusion is not justified since nothing shows that SMC4 is phosphorylated by Plk1 at the mutated sites and nothing even shows that SMC421A restricts chromosome condensation. Even if these points were demonstrated, there would still be the need to show that the regulation of condensation by Plk1 is the mechanism which transcription is regulated (point 1).

*We revised our wording to temper the statement.*

5. Further questioning the model proposed, the literature shows that Plk1, by phosphorylating condensin, promotes chromosome condensation in mitosis rather than restraining it.

*We discussed the effect of other Plk1-phosphorylated condensin II components in the text. We observed an opposite effect of Plk1 activity, likely due to the different mitotic stages examined. Plk1 activity in prophase phosphorylates condensin II components to establish condensation, while in prometaphase, we propose it restrains condensation.*

6. As part of the investigation of a potential role of Plk1 in retraining condensation, it would be relevant to visualize and quantify condensin levels on chromosomes.

*In Fig. 5A (previously Fig. 4A), we showed the chromosomal-bound SMC4 protein level in mitotic cells indicated by the Halo tag. SMC4 levels increased after Plk1 inhibition, suggesting an increase in condensin levels.*

Minor points:

7. The authors seem to propose that the actual process of transcription, rather than some of the gene products, is required for normal chromosome segregation. In the absence of a good way to discriminate between them, both possibilities should probably be mentioned and discussed.

*We mentioned this in the discussion section.*

8. Line 233-234: "In these cells, Plk1 activity is sufficient for cohesin removal during prophase but insufficient to maintain proper chromosome condensation."

This should be rephrased since it is more a question of timing of Plk1 inhibition rather than a question of different Plk1 activity levels. Cells were simply arrested in prometaphase (after prophase) with STLC before inhibiting Plk1.

*We rephrased the sentence.*

9. In Fig 2A, it would be good to specify which treatment(s) the 2 cells shown are taken from.

*We added the information in the figure legend.*

10. Some of the statistical test results are surprising in view of the data shown in the figures. For example, in Fig 2H, it is hard to believe that the p-value is above 0.05 when comparing Flag and Flag-Plk1WT after DMSO treatment. Could it be that the Plk1AS is not fully functional?

*We checked the statistical analysis results presented in Fig. 2H. A two-way ANOVA was conducted on the entire dataset, followed by Tukey's test for multiple comparisons. The adjusted p-value for the comparison between FLAG and FLAG-Plk1^WT^ groups was 0.2114.*

11. In Fig 3J, I presume that the peaks shown correspond to only those shown in Fig 3I rather than all peaks in the entire genome? This should be clarified. In Fig 3K, label the X axis as "Log2 Fold Change Plk1 inhibition / control". And define "padj".

*The peak overlay shown in Fig. 3I (now Fig. 4C) includes all mapped peaks. We mentioned this in the figure legend. We modified the label and legend of Fig. 3K (now Fig. 4E).*

12. As a piece of advice for the future, the visual rendering of the model shown in Fig 4G could be improved to show more clearly the implication of Plk1, SMC4 and condensins. However, I am not suggesting making a new version until the mechanisms are more clearly defined based on experimental evidence.

*We modified the model figure.*

**Reviewer 3:**

Major points:

1. Throughout the manuscript the authors rely on an EU incorporation assay to measure transcription during mitosis, but the details of the assay and quantitation are not properly documented and the results are in contrast to many studies in the literature. Specific concerns are listed below:
   a. In Figure 1AB the authors label all transcripts in cells treated with STLC and with or without Plk1 inhibition. In this experiment they do not distinguish between mitotic transcription and transcription that occurs in G2. This needs to be clarified by only examining transcription in mitotic cells (as presented in Figure 1DE).

Furthermore ~75% of all cellular transcription is rRNA production. The magnitude of EU incorporation reduction suggests that Plk1 inhibition has a strong effect on rRNA transcription. This should be examined and discussed.

*We discussed the effect of G2 transcripts in the 5-EU experiment in the text. We specifically tested the effect of Plk1 activity on mitotic and G2 transcripts in Figures 1C-E. RNAPII is the major polymerase function during mitosis. According to PMID: 26527278, rRNA production is not a major component of mitotic nascent transcripts. We don’t think Plk1 regulates RNAP I.*

b. In Figure 1DE the authors examine EU labeling of mitotic chromosomes in control and BI-treated cells. In all samples they find that EU labels the entire chromosome in a pattern indistinguishable from DAPI. This is in stark contrast to many studies in the literature (e.g. 29626011, 26190260, 28760200, 32035037, 28912132, 22308327) that show EU labeling and RNAPII predominately at the centromeres of mitotic chromosomes. The authors need to explain why their results are dramatically different than many other published studies. Additionally, the authors need to include a no EU control to ensure that the signal they observed comes from nascent transcripts. Their results would be solidified if they could provide an orthogonal validation of the EU labeling experiments. Appropriate methods could include RNAPII pS2 staining or EU-labeling followed by RNA sequencing.

*We included a no 5-EU control (Figure S1A). We did not observe an enrichment of 5-EU at centromeres, likely due to the process of chromosome spreading. While we did not find other studies performing 5-EU staining, PMID: 33881484 evaluated 5-EU-incorporated centromeric transcripts by RT-qPCR, confirming that centromeric transcripts were labeled.*

c. In Figure 2 the authors have the perfect control for their experiments. The authors should arrest RNAPII-AID cells in mitosis, degrade RNAPII, then label with EU. This will provide a baseline for the maximal amount of reduction that could be expected with complete transcriptional inhibition.

*We acknowledge that synchronizing POLR2A-AID cells and depleting RNAP II in mitosis would serve as a great control for the 5-EU incorporation experiment. However, due to technical difficulties, we were unable to achieve good synchronization in POLR2A-AID cells.*

d. In the panels in this figure and all figures the authors have not included the necessary information about how image quantitation was performed and what statistical tests were used to compare samples (this information is not present in the methods or figure legends). In the current form it is impossible to evaluate the data presented in this manuscript. In addition, the meaning of the * symbols on the figures are not indicated. This is of particular concern because small differences (e.g. Figure 1I, ~10% difference between control and treated) lead to apparently extremely significant p-values. I suspect that the authors are treating each cell as a n instead of each biological replicate as a n. I would recommend that the authors consult 32346721 for appropriate use of statistics in cell biology experiments.

*We updated the description of the statistical analysis in the methods section.*

2. This manuscript focuses on the role of Plk1 in regulation of mitotic transcription but largely ignores important literature related to this topic. Plk1 is a critical regulator of the 'prophase pathway' of cleavage-independent removal of cohesin from mitotic chromosomes (11931760 and many others). Additionally, a recent study showed that removal of cohesin from chromosomes by the prophase pathway and WAPL is important for mitotic transcriptional inhibition and retention of elongating RNAPII at the centromere (32035037). In fact, 32035037 showed that inhibition of Plk1 using BI2536 leads to an increase in RNAPII on mitotic chromosomes in human RPE1 cells and Xenopus egg extracts (Figure S3). This contrasts with the statement on line 376. These results are in direct contrast to the results presented in the current manuscript. The authors need to discuss the relationship of their findings to this previous work and explain why they arrive at a completely different conclusion.

*We discussed the effect of other Plk1’s functions in the text. We observed an opposite effect of Plk1 activity, likely due to the different mitotic stages examined. Plk1 activity in prophase promotes cohesin removal and RNAP clearance, while in prometaphase, we propose it retains RNAP levels on chromosomes, likely by maintaining chromosomal accessibility.*

3. In Figure 2A the authors treat asynchronous cells for 5 hours with alpha-amanitin and observe mitotic defects. They conclude that mitotic transcription is important for normal mitosis. However, they cannot make this conclusion form their experimental design. The time window of treatment will lead to transcriptional changes in G2 and mitosis. Transcriptional changes in G2 could have an indirect effect on mitotic progression. The authors could control for this effect by collecting cells in mitosis by shake-off and then inhibiting transcription. Alternatively they could use fast-acting RNAPII inhibitors (e.g. triptolide, cdk7 or cdk9 inhibitors). Additionally, the results presented here contrast with studies in model organisms (Drosophila, Xenopus, Zebrafish) showing that transcriptional inhibition during early development does not cause mitotic defects (e.g. 35015118). The authors need to address this literature and the issues with their experimental design before these experiments will be interpretable. These timing concerns also concern RNAPII-AID experiments as well.

*We agree that controlling the effect of G2 transcripts is important. However, we were unable to achieve good synchronization in POLR2A-AID cells. PMID 30080136 suggested that triptolide, but not α-amanitin, efficiently inhibits transcription, but the α-amanitin concentration they used may be too low for effective inhibition (PMID: 38276537). We have tried inhibiting transcription using triptolide, and it showed minor effect on mitotic transcription and fidelity. Triptolide is implied to have complicated effects on cells other than transcription inhibition (PMID: 38293343). Actinomycin D significantly decreased mitotic transcription, but it is likely through DNA-intercalating function. (1 biological replicates performed in EU assay, 3 biological replicates performed to assess mitotic error rate. Both assays were done in RPE1 Plk1^AS^ cells.)*

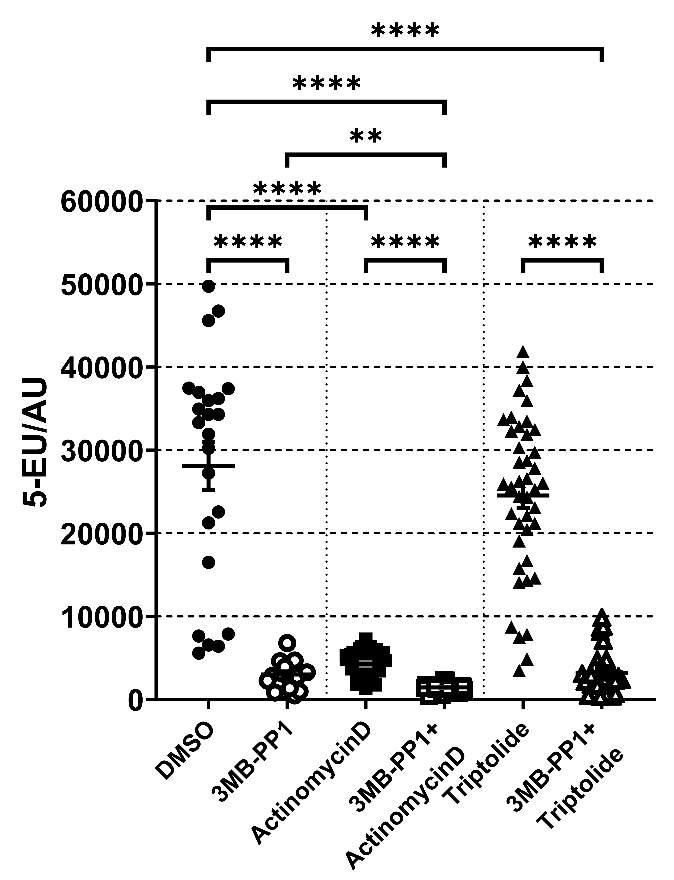

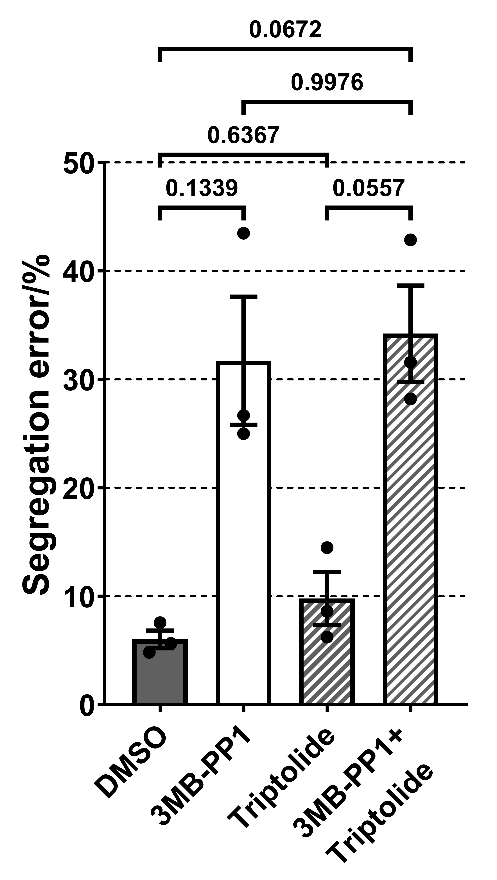

4. In Figure F-H the authors should provide western blots showing expression levels of rescue constructs relative to endogenous Plk1. Additionally, authors should provide microscopy images verifying expected localization of indicated constructs.

*The cell lines used in Fig. 2F-H were reported and validated in PMID: 31468671. Localization of FLAG-Plk1^ΔC^-H2B is shown in Fig. S2B.*

5. Figure 3. The authors should provide a Venn diagram of the number of peaks and peak overlap in ATAC-seq experiment between all samples. In Figure 3K it is not clear what is being tested here. The information in the figure legend is not sufficient to evaluate. If the authors want to preform differential peak analysis they should merge all peaks from all replicates and then calculate read differences between different experimental samples. One example can be found here: 30361341. In Figure 3J the authors should use Deeptools to provide a metaplot of all samples using annotated gene models to support their hypothesis that Plk1 controls access to promoter regions.

*We included the Venn diagram (Fig. 4D and Fig. S4B) and Deeptool profile (Fig. 4C and Fig. S4D). We modified our analyses.*

6. In Figure 4 the authors introduce an AID-Smc4 but do not use it to test their hypothesis. They propose that inhibition of Plk1 leads to more condensin on chromosomes which results in less mitotic transcription. If this idea is correct then depletion of Smc4 should rescue EU incorporation in BI-treated cells. This is a critical experiment to test their hypothesis.

*We agreed that the combination of condensin depletion and Plk1 inhibition is a great experiment to test our hypothesis. We tried the treatment (STLC+5-EU+BI2536+PhIAA). However, the timeline for SMC4 degradation is approximately 2 hours, resulting in a heterogeneous population of prometaphase cells with varying levels of chromosome condensation. The rapid reduction of Plk1 activity induced by BI2536 further contributed to data variability,* *making it too noisy to analyze.*

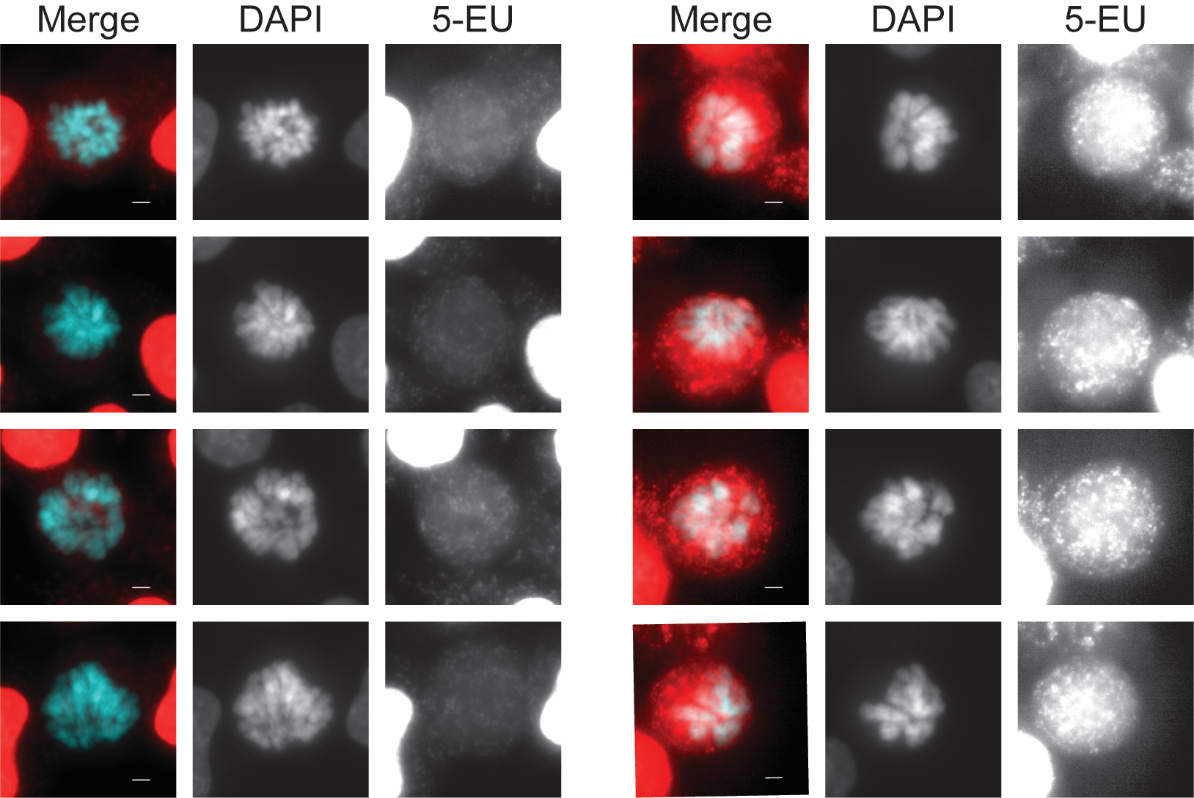

7. The authors should show evidence of reduced phosphorylation of Smc4-21A by changes in mobility by western blot. They should also show the localization of this construct by microscopy.

*We included a western blot (Fig. S5A, original gel image below “#2 wt” and “#7 allA”) to show mCherry-SMC4^WT^ and SMC4^21A^ expression. However, we did not observe a phosphorylation shift in SMC4^21A^ compared to SMC4^WT^. It is possible that not all 21 sites in SMC4^WT^ are phosphorylated in the cell, which could result in only a slight change in mobility. Additionally, not all phosphorylation events would cause a protein mobility change.*

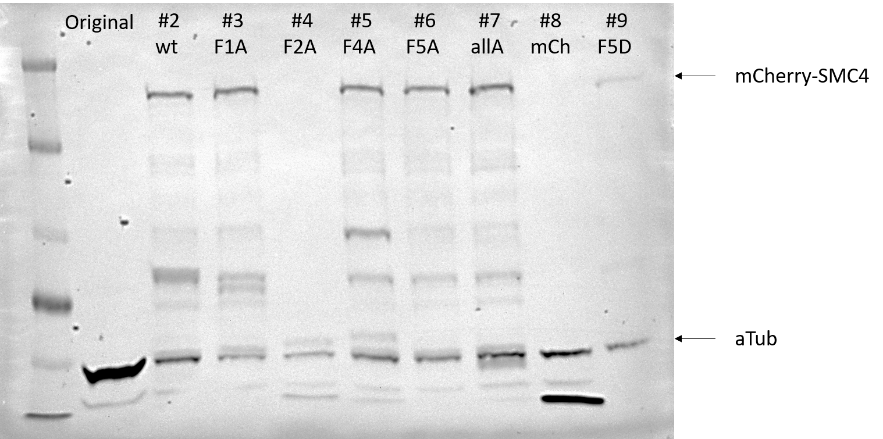

*We included immunofluorescent images (Fig. S5B-C) to show mCherry-SMC4^WT^ and SMC4^21A^ localization and expression.*

8. Line 297. Authors state that Smc4-AID and Smc4-21A cells are viable, but the data is not present in the manuscript. This statement is in contrast to the Broad DepMap which shows that Smc4 is a common essential gene required for cell growth of 1149 cancer cell lines.

*We included the growth assay result (Fig. S5D) to show the growth of mCherry-SMC4^WT^ and SMC4^21A^ cells. SMC4-AID cells are not viable with PhIAA depleting SMC4, confirming that SMC4 is an essential gene.*

Minor points:

There are a number of phrasing errors that need to be corrects:

a. Line 214 "Chromatin condensed into chromosome starts from early prophase."
b. Line 345 "The underlying mechanisms regulating chromosomal remain poorly understood."
c. Line 363 "Interestingly, blocking Plk1 phosphorylation of CAP-D3 and CAP-H2 resulted in the retention of chromosome condensation"

*We made changes to the sentences you mentioned.*
